# Predicting recognition between T cell receptors and epitopes using contextualized motifs

**DOI:** 10.1101/2022.05.23.493034

**Authors:** Emmi Jokinen, Alexandru Dumitrescu, Jani Huuhtanen, Vladimir Gligorijević, Satu Mustjoki, Richard Bonneau, Markus Heinonen, Harri Lähdesmäki

## Abstract

We introduce TCRconv, a deep learning model for predicting recognition between T-cell receptors and epitopes. TCRconv uses a deep protein language model and convolutions to extract contextualized motifs and provides state-of-the-art TCR-epitope prediction accuracy. Using TCR repertoires from COVID-19 patients, we demonstrate that TCRconv can provide insight into T-cell dynamics and phenotypes during the disease.

## Main

T-cell receptors (TCRs) form diverse repertoires through V(D)J recombination, which allows T-cells to recognize a large variety of antigens. Short peptide sequences from the antigens, called epitopes, are presented to T-cells via major histocompatibility complex (MHC) molecules, and successful recognition of an epitope-MHC complex by a TCR results in T-cell activation. Discovering epitope-specific TCRs holds the potential to provide clinically relevant insights into TCR repertoires in fields ranging from vaccine design and diagnostics to immunotherapy biomarker identification.

Latest high-throughput sequencing technologies have enabled profiling large quantities of TCR sequences. Concurrently, several methods have been proposed for predicting TCR-epitope recognition ^1,2,3,4,5^. Previous work has shown that while the complementarity-determining region 3 (CDR3) is crucial for the prediction, it is beneficial to utilize also other TCR regions, and the paired TCR*αβ* sequences ^1,2,3,4^. We focus on major open questions in TCR-epitope prediction: how to utilize efficiently all TCR regions that determine epitope-specificity, handle TCR cross-reactivity, and use TCR-epitope prediction methods for unsupervised analysis of TCR-repertoires.

Here, we present TCRconv, a convolutional deep neural network that utilizes rich contextualized transformer embeddings of TCRs to predict epitope recognition (see Fig. 1a, Supplementary Fig 1, and Methods for details). Unlike the previous methods, TCRconv models TCR specificity as multilabel predictor that naturally accounts for TCR cross-reactivity. The transformer model BERT (Bidirectional Encoder Representations from Transformers) transfers information from the complete TCR sequence to the CDR3 embedding from which the convolutional networks then extract and utilize contextualized motifs.

**Figure 1:**
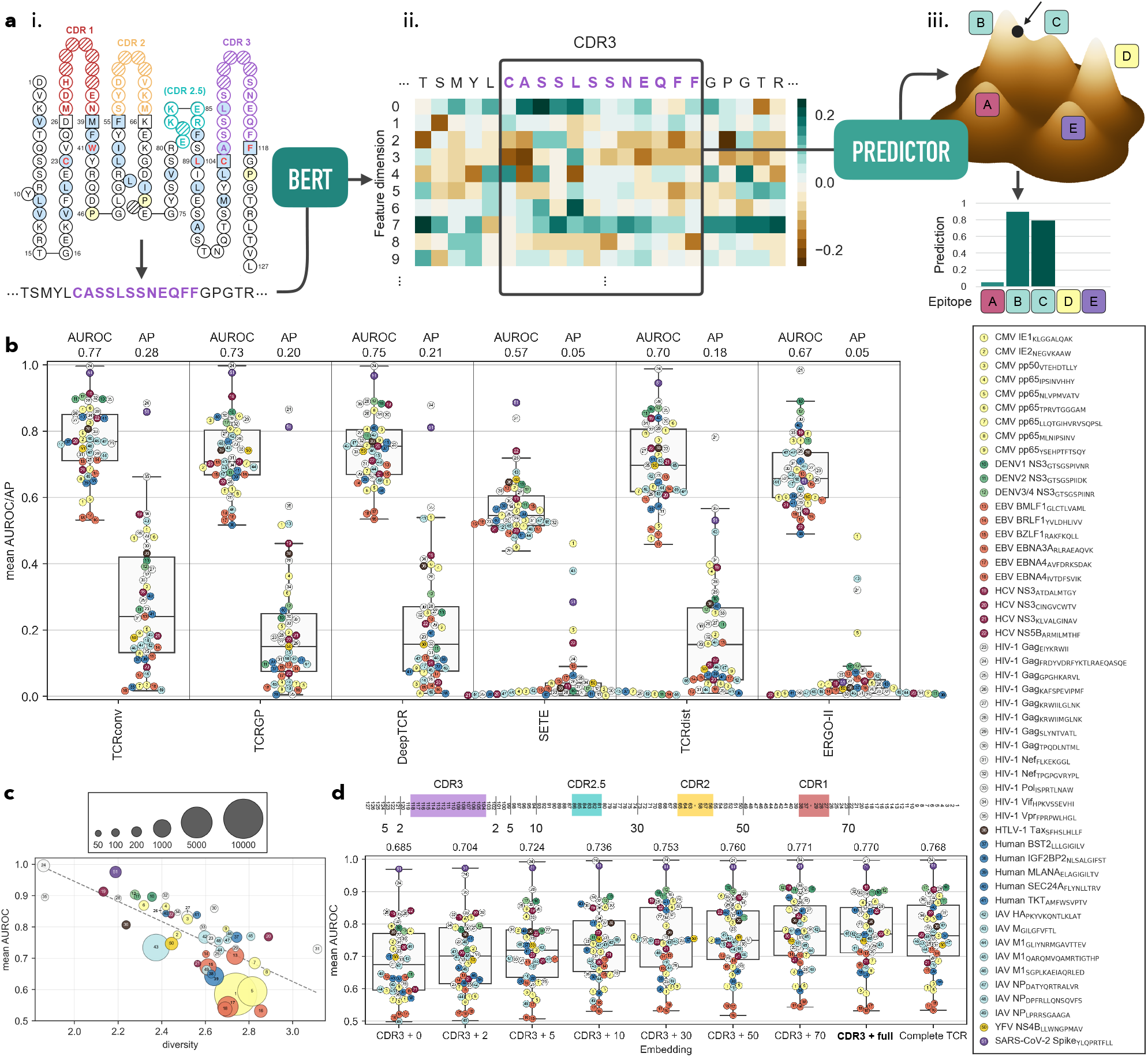
TCRconv evaluation. (a) TCRconv pipeline. (i) The TCR sequence determined by V(D)J recombination contains the complementarity-determining regions. TCR*α* and/or TCR*β* sequences can be used, here TCR*β* is shown. (ii) ProtBERT embedding is created for each TCR sequence and the CDR3 embedding, transfused with information from its context, is extracted. (iii) The multilabel predictor produces simultaneously separate predictions for each epitope. (b) Comparing TCRconv to other methods using average AUROC and AP scores. (c) The AUROC scores for TCRconv predictions correlate negatively with the diversity of the epitope specific TCRs (Pearson correlation −0.72). (d) Increasing the embedding context size increases the predictive AUROC score. The schematics on top show the approximate sections included in different context sizes. CDR3 + X refers to CDR3 embeddings with context size X and complete TCR to embeddings for complete TCRs without extracting only the CDR3 parts. TCRconv uses CDR3 + full (bolded). Results for panels b-d are obtained using stratified 10-fold cross-validation on VDJdb*β*-large dataset.

We compared the accuracies of TCRconv and previous methods that include Gaussian processes (TCRGP^2^), deep learning methods (DeepTCR^3^, ERGO-II^4^, SETE^5^), and TCRdist^1^. We collected two epitope-specific TCR*β* datasets from VDJdb^6^, a comprehensive VDJdb*β*-large consisting of data with all confidence levels and a smaller high-quality VDJdb*β*-small (Supplementary Table 1 and Methods). Prediction accuracies are quantified using average precision (AP), which accounts for class imbalances, and area under the receiver operating characteristic curve (AUROC). TCRconv achieves the highest AP and AUROC scores on VDJdb*β*-large (33% and 3% improvement to the second best DeepTCR) (Fig. 1b). High AP scores are essential as minimizing false positive predictions with large TCR repertoires and small TCR clones^7^ is crucial. Overall, all methods performed better on the higher confidence, albeit smaller, dataset VDJdb*β*-small (Supplementary Fig. 2, Supplementary Table 2).

As TCRs can be cross-reactive (Supplementary Fig. 3), TCRconv benefits from using a single multilabel predictor that can predict a TCR to recognize several epitopes. Whereas, to account for cross-reactivity with previous binary^1,2^ and multiclass classifiers^3,5^, a large set of separate (one-vs-all) classifiers had to be trained, one for each epitope. TCRconv performs well also with cross-reacting TCRs (Supplementary Fig. 4). Further, consistent with previous results1,2, we confirmed that prediction accuracy across epitopes correlates negatively with the diversity of the TCRs recognizing these epitopes (Fig. 1c, Supplementary Fig. 5a, and Methods).

CDR3 is essential in epitope recognition, but structural^8^ and computational^1,2^ evidence suggests that CDR1 and CDR2, that mainly contact the MHC, may also interact with the epitope and aid the prediction. We next evaluated how much of the TCR sequence around the CDR3 should be used as context. The prediction AUROC score improves gradually from 0.68 to 0.77 when the context size is increased from no context to full context on VDJdb*β*-large (Fig. 1c), indicating that BERT successfully conveys relevant information from the context to the CDR3 embedding. With both datasets the AUROC and AP scores improve or remain the same when using context before CDR1 (Supplementary Fig. 5b). Remarkably, the entire TCR embedding is not needed, but using the CDR3 embedding with full context provides similar or slightly better results. The MHC can also affect the TCR-epitope recognition, but our analysis suggests the HLA-types do not have a substantial effect on the prediction (Methods and Supplementary Fig. 6).

We studied the effect of TCR*α* and TCR*β* on TCR-epitope prediction on VDJdb*αβ*-large dataset of paired TCR*αβ* sequences (Supplementary Table 1 and Methods). We find substantial performance improvement from using both chains over either chain individually (Supplementary Fig. 7).

Finally, we demonstrate how to utilize TCRconv on repertoire data to track T-cell dynamics^9^ during COVID-19 and reveal the phenotypes of SARS-CoV-2 specific T-cells in moderate and severe COVID-19. We first trained a TCRconv model specifically for SARS-CoV-2 epitopes using ImmuneCODE^10^ and VDJdb*β*-large data (Methods, Supplementary Fig. 8, and Supplementary Table 3).

To track the T-cell dynamics, we predicted each TCR’s specificity to the selected SARS-CoV-2 epitopes in 110 healthy control^11^ and 493 COVID-19 patient^10^ repertoires (Supplementary Table 4). For each repertoire, we quantified the normalized frequency of TCRs predicted to be SARS-CoV-2-specific during COVID-19. Fig. 2a shows that the frequency of SARS-CoV-2-specific T-cells is highest during the first two days after diagnosis and starts to decrease during the first week. In contrast, with IAV, CMV, EBV, and HCV (Fig. 2a and Supplementary Fig. 9) the normalized frequency remains lower.

**Figure 2:**
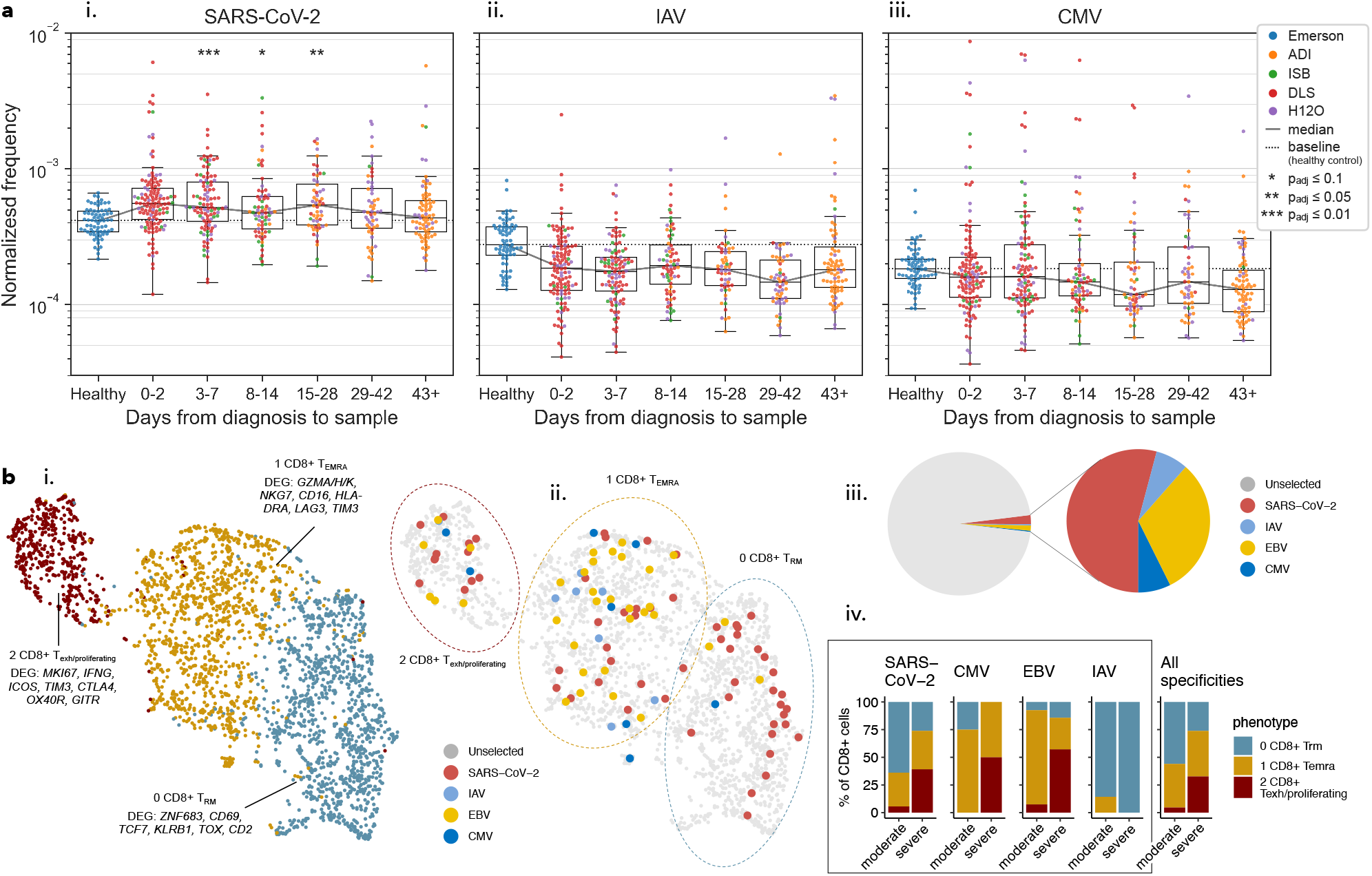
Analyzing TCR repertoires of COVID-19 patients with TCRconv. (a) Dynamics of (i) SARS-CoV-2 (ii) IAV, and (iii) CMV-specific T-cells in terms of frequency normalized by the number of virus-related epitopes. There are 20 epitopes for SARS-CoV-2, eight for IAV, and nine for CMV. Each data point corresponds to a repertoire and is colored by its dataset (Supplementary Table 2). Symbols “*” indicate statistically significant increase in frequency compared to Healthy samples (see Methods and Supplementary Table 5). (b) Phenotypes of SARS-CoV-2-specific CD8+ T-cells from bronchoalveolar lavage samples from patients with moderate (n=3) or severe (n=6) COVID-19 disease. (i) UMAP representation of CD8+ T-cell phenotypes, (ii) clustering with epitope-specific T-cells marked, (iii) proportions of epitope specific T-cells, (iv) phenotype distribution of virus specific T-cells.

To link TCR-specificity to T-cell phenotype, we utilized scRNA+TCR*αβ*-seq of CD8+ T-cells from bronchoalveolar lavage samples of nine COVID-19 patients^12^. As expected, SARS-CoV-2-specific T-cells were more abundant than T-cells recognizing other tested viruses (CMV, EBV, IAV) (Fig. 2b). Moreover, in patients with moderate disease (n=3) the SARS-CoV-2-specific T-cells most often had tissue-resident memory phenotype (overexpression of ZNF683, CD69, TCF7) (Fig. 2b, Supplementary Fig. 9). In patients with severe disease (n=6), we found SARS-CoV-2-specific T-cells to have possibly overtly proliferating (MKI67) and exhausted (HAVCR2/TIM3, CTLA4) phenotype, with high expression of co-stimulatory signals (ICOS, TNFRSF4/OX40R, GITR) and IFNG (Fig. 2b). These findings refine previous findings^13^ by suggesting that patients with a moderate disease course form T-cells capable of elimating SARS-CoV-2 with minimal tissue damage while T-cell overactivation in patients with a severe disease leads to an inappropriate tissue damage.

TCRconv is available at github.com/emmijokinen/tcrconv.

## Methods

### Data

For training and testing our model, we have constructed three data sets of human TCR-sequences from the data available in the VDJdb database^6^ (vdjdb.cdr3.net). VDJdb gives confidence scores from 0-3 for each of its entries, 0 for low confidence or no information (a critical aspect of sequencing or specificity validation is missing), 1 for moderate confidence (no verification or poor TCR sequence confidence), 2 for high confidence (has some specificity verification, good TCR sequence confidence), and 3 for very high confidence (has extensive verification or structural data), see more detailed description in github.com/antigenomics/vdjdb-db. For VDJdb*β*-large, we selected TCRβs with all confidence scores, and with at least 50 unique TCRβs for each epitope. This resulted in a data set with 51 distinct epitopes and 30 503 unique TCRβs. For VDJdb*β*-small we chose TCRβs with at least a confidence score of 1 and at least 40 unique TCRs per epitope, which resulted in 1977 unique TCRs specific to 21 epitopes. Finally, VDJdb*αβ*-large consists paired TCR*αβ* sequences with all confidence scores and at least 50 unique TCR*αβ*s per epitope, in total 20200 unique TCRs and 18 epitopes. A TCR is considered as unique if the combination of its CDR3 and V- and J-genes is unique. Table 1 summarizes these datasets and the cross-reactivities of the TCRs are visualized in Supplementary Fig. 3. As the requirements for the datasets overlap, so do the datasets, for example VDJdb*β*-large contains the complete VDJdb*β*-small dataset. VDJdb*β*-large and VDJdb*β*-small were collected in January 2021 and VDJdb*αβ*-large in September 2021 which explains why some of the SARS-CoV-2 epitopes are only present in VDJdb*αβ*-large. All presented model evaluations are conducted using a stratified 10-fold cross-validation, where TCRs specific to each epitope are distributed to the folds as evenly as possible. As our data set only consists of unique TCRs, the same TCR can never be both in training and test folds.

### TCR Embeddings

Transformer based language models for proteins, such as BERT (Bidirectional Encoder Representations from Transformers), can capture protein folding as well as learn useful representations of binding sites and complex biophysical properties^14^. They have been successfully used in various tasks, including protein family and protein interaction prediction^15^ and protein-specific drug generation^16^, making them a plausible choice for modeling TCRs as well. For constructing the TCR embeddings we used ProtBERT^17^ which is trained on 216 million UniRef100 sequences. The model was trained with a token-prediction task and during the training phase 15% of the tokens (amino acids) in the sequences were replaced by a MASK token. The model contains 16 attention heads in each multi-head attention block on 30 layers, with 420 million parameters in total. The embedding dimension for each amino acid in a sequence is 1024, resulting in an embedding dimension of *L ×* 1024 for a sequence of length *L.* We constructed the embeddings for TCR*β*-sequences, defined by V**β*-* genes, CDR3*β* sequences, and J*β*-genes, whose lengths varied between 103-137 amino acids. For our final embeddings, we extracted only the sections corresponding to the CDR3*β* sequences from the TCR*β*-embeddings.

We also attempted to make the ProtBERT model more specialized to TCR sequences by fine-tuning it on 5 million TCR*β* sequences from VDJdb^6^, and studies of Emerson et al. 11 and Dash et al. 1 for 8 epochs but this did not improve the prediction accuracies (mean AU-ROC 0.848 and AP 0.575 on VDJdb*β*-small dataset). We also tested two ELMo (Embeddings from Language Models) architectures, classical ELMo^18^ and masked ELMo^19^, and trained them on a smaller dataset of 3 million TCR*β*-sequences from the same sources as those used in the BERT fine-tuning. The main difference between these two models is that instead of unidirectional LSTMs, the masked ELMo uses a bidirectional two-layered LSTM and when trained in the token prediction task, the predicted token (amino acid) is masked to avoid leakage of information. We found that both ELMo models produced reasonable accuracies in the prediction task (mean AUROC and AP 0.838 and 0.539 for ELMo and 0.847 and 0.571 for masked ELMo, on VDJdb*β*-small dataset), and with masked ELMo we achieved almost as good accuracy as with the BERT embeddings.

### CNN classifier

Our multilabel classifier consists of a parallel convolutional unit and a simple linear unit. The classifier was modified from the CNN classifier presented by Glig-orijević et al.^20^ As shown in Supplementary Fig. 1, the convolutional unit consists of parallel convolutional layers with varying kernel sizes (5, 9, 15, and 21, with 120, 100, 80, and 60 filters, respectively) that can capture different length motifs. The outputs from these layers are concatenated and fed through batch normalization, rectified linear unit (ReLU) activation, and a dropout layer with 0.1 dropout. Those are followed by another convolutional layer (kernel size 3, 60 filters) that can extract higher level features based on the outputs from the previous convolutional layers. Finally, max pooling is performed over the sequence lengths, which provides fixed sized outputs regardless of the sequences’ lengths. The linear unit can more flexibly utilize the expressive features of the BERT embeddings. It consists of a max pooling layer, a linear layer, and a ReLU activation. The outputs of the convolutional and linear units are concatenated and put through a dropout layer with dropout 0.1 and batch normalization and ReLU. The final linear layer gives predictions simultaneously for each class that are separately squashed between 0 and 1 by a sigmoid layer.

For optimizing the network parameters, we use binary cross-entropy which allows predictions for multiple epitopes to be 1. Positive answers for class c are weighted by *p_c_* = *n*_TCRs_(|*c*| · *n*_classes_), where *n*_TCRS_ is the number of TCRs in the training data, |*c*| is the size of class *c*, and *n*_classes_ is the number of classes. We use a learning rate of 0.0002, except for the linear unit, for which the learning rate is set to of 0.01.

For training the models, we use stochastic weight averaging (SWA) with learning rate scheduling^21^. The models are first trained for 2500 iterations (minibatches) without weight averaging but with cosine annealing for the learning rate, so that the learning rates gradually decrease. After that, the training is continued for another 500 iterations with stochastic weight averaging on every iteration and again a decreasing learning rate is used.

### Comparison to other methods

We compared TCRconv to recently published methods for predicting TCR epitope-specificities, TCRGP^2^, DeepTCR^3^, SETE^5^, TCRdist^1^, and ERGO-II^4^. Apart from ERGO-II all the methods used epitopes as class information to predict if a TCR would recognize one of the predetermined epitopes. TCRGP is a Gaussian process-based classifier that can utilize CDR3 or additionally any or all of the other CDRs from either *α*- or *β*-chain or both, depending on what information is available. Similarly to TCRconv, DeepTCR uses convolutional neural networks, but with trainable embedding layers for the CDR3s and V/D/J genes. SETE on the other hand takes a PCA of 3-mer occurrences in CDR3s and uses gradient boosting decision trees to classify CDR3-sequences, and TCRdist uses a BLOSUM62 based distance measure between selected CDRs to determine if new TCRs are closer to TCRs specific to a certain epitope or to some control TCRs. ERGO-II learns LSTM encodings for both CDR3s and epitopes (or autoencoder embeddings for CDR3s) and uses both the TCR and the epitope as inputs to predict if a given TCR and epitope bind.

With TCRGP, DeepTCR, SETE, and TCRdist we trained separate binary classifiers for each epitope, so that TCRs known to recognize the epitope in question are considered as positive data points and TCRs specific to other epitopes are considered as negative data points. DeepTCR and SETE had options for multiclass classification, but they did not provide support for cross-reactive TCRs that our data contains. Therefore, they would have had a disadvantage if trained as multiclass classifiers as they then would have operated with either conflicted or missing class labels.

We compared the above methods on our two data sets using stratified 10-fold cross-validation. The TCRconv model was trained once for each fold and all the known epitope-specificities are determined by multi-hot encodings. With TCRGP, DeepTCR, SETE, and TCRdist a separate model was trained for each epitope in each fold. The folds used in the cross-validation were the same for each of these methods. As suggested by the authors, with DeepTCR 25 % and with ERGO-II 20 % of the training data was used as validation data for determining early stopping when training the classifiers. With ERGO-II, also as suggested by the authors, we used all the positive TCR-epitope pairs in our data, but additionally sampled five times more negative data. Therefore, when with TCRconv e.g. TCR CASLS-GRAPQHF, TRBV27*01 occurs once in the data set with a multi-hot encoding indicating that it can recognize epitopes GTSGPIINR and GTSGPIVNR, with ERGO-II it is repeated 12 times, twice in a positive pair with both GTSGPIINR and GTSGPIVNR, and ten times in negative pairs formed by randomly selecting ten of the other 19 epitopes in the VDJdb*β*-large data set.

### TCR diversity

To estimate the diversity of *N* TCRs specific to a certain epitope, we utilized a diversity measure similar to measures used in previous studies^2,7^. These measures are based on Simpson’s diversity index, but due to the large variety of TCRs they measure similarities between TCRs instead of exact matches. Here the similarity between TCRs *i* and *j* is computed based on the used embeddings x*_i_* and x*_j_* that have been aligned based on IMGT numbering:

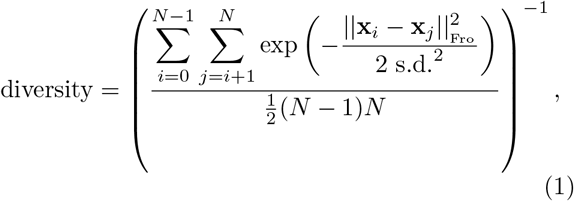

where s.d. is set to 10.4 (maximum feature-wise standard deviations multiplied by the median sequence length 14.

### The effect of HLA-type on the predictor

As the TCR regions outside the CDR3 do not often interact with an epitope but may interact with the MHC molecule presenting the epitope, the HLA-type of the MHC can affect the recognition between the TCR and the epitope. Therefore, utilizing these regions could introduce a bias in the epitope-specificity prediction. Although the available TCR-epitope-MHC complexes in VDJdb6 contain various HLA-types, most of the data is restricted to HLA-A*02 and almost all the epitopes are presented by a single HLA-group (see Supplementary Fig. 6a). This makes it difficult to model or even assess how different HLA-types affect the TCRs’ ability to bind certain epitopes. To ensure that our multilabel predictor is predicting a TCR’s ability to bind to an epitope and not to the HLA presenting it, we examined how much the results differ between different HLA-genes, and if TCRconv is trained on data restricted by any HLA type or only with HLA-A*02 restricted data. The number of epitopes restricted by each HLA-gene is limited and the prediction accuracy varies considerably between epitopes, but Supplementary Fig. 6 indicates that differences between HLA-genes are modest (AUROC across genes varies from 0.743 to 0.810 while AUROC across epitopes varies from 0.532 to 0.996); panel b) and that the accuracy is similar when TCRconv is trained on all data or only with HLA-A*02 restricted epitopes (panel c). These results suggest that TCRconv predicts TCR’s ability to bind epitopes and not the HLAs.

### TCRconv model for SARS-CoV-2 epitopes

For training TCRconv models for SARS-CoV-2 specific epitopes, we utilized ImmuneCODE10 MIRA-data of TCRs specific to MHC-I restricted peptides. To fully exploit the data, we did the following preprocessing with three options for the TCR-sequences: 1) If the V- and J-genes and their alleles could be determined from the nucleotide sequence (length 29 nucleotides), we used the exact TCR amino acid sequence determined by the CDR3*β*, V- and J-genes. 2) If a V- or J-gene could be determined but not its allele, we set the allele to 01 and used it for constructing the amino acid sequence. 3) If a gene could not be determined, we utilized a partial amino acid sequence that we could determine based on the nucleotide sequence. BERT embeddings were computed for these TCR*β* sequences and the parts of the embeddings corresponding to the CDR3s were extracted and used with the classifier. TCR uniqueness was determined by these longest amino acid sequences we could obtain. We selected 139099 unique TCRβs specific to 188 peptide groups with at least 50 unique TCRβs specific to them (see Supplementary Table 3) and used stratified 10-fold cross-validation with these TCRs to train TCRconv model, whose performance in terms of mean AUROC and AP is shown in Supplementary Fig. 8a. Supplementary Fig. 8c shows AUROC and AP scores, and diversity of TCRs specific to each peptide group by their genomic location. We then selected the twenty peptide groups that performed best in terms of weighted mean of AUROC and AP scores (both scores were scaled into range [0,1]) and used the TCRs specific to them with VDJdb*β*-large dataset to construct the final predictor (performance using stratified 10-fold cross-validation is shown in Supplementary Fig. 8b).

### Frequency of SARS-CoV-2 specific TCRs in repertoires

We utilized TCR repertoires from ImmuneCODE that contain at least 250000 TCRs and the number of days between diagnosing the patient and collecting the sample is reported. As control data we used TCR repertoires of healthy subjects from Emerson et. al. 11 that also had at least 250000 TCRs and where the subject age is at least 18 years. The data is described in Supplementary Table 4. The sequences were preprocessed in the same way as the ImmuneCODE MIRA-data and each sample was downsampled to 250000 TCRs. Using the TCRconv model for SARS-CoV-2 epitopes, we predicted the specificity of each T-cell within these repertoires. We chose a threshold separately for each epitope that corresponds to false positive rate of 0.001. With thresholds this strict we are not likely to find all TCRs specific to the selected epitopes but have a high confidence in that the TCRs predicted to recognize these epitopes are true positives. We computed the frequency of TCRs predicted to be specific to the SARS-CoV-2 and normalized it by the number of SARS-CoV-2 epitopes (20) to be better able to compare to responses for other viruses. A Linear regression analysis was performed to assess if COVID patients have significantly higher frequency of virus specific T-cells than healthy control subjects, and if the frequencies are positively correlated with subjects’ age. This was done separately for each time interval using linear model *y* = *a* + *b_cc_x_cc_* + *b_age_x_age_*, where *y* is frequency, *a* is offset, *b_cc_* is parameter for case-control covariate *x_cc_* which is zero for control samples and one for case samples of the considered time interval, and *b_age_* is parameter for age covariate *x_age_*. See Supplementary Table 5 for significance of the parameters for case-control and age covariates measured by one-tailed t-test.

### Phenotypes of SARS-CoV-2 specific T-cells in moderate and severe COVID-19

Count matrices, TCR*αβ*-seq results, and metadata from Liao et al12 were downloaded from GEO GSE145926. The data was analyzed mainly with Python package scVI tools^22^ (v 0.14.5) and R package Seurat^23^ (v 4.0.4). Cells with > 10 % mitochondrial gene counts, < 1000 UMI counts, < 200 or > 6000 detected genes, and cells with no detected TCR were filtered out. The highly variable genes were identified with “highly_variable_genes” function in scVI tools with default parameters, which were then used to learn latent embeddings with “model.SCVI” function in scVI tools with default parameters. The CD8+ T-cells were then identified with SingleR^24^ (v 1.6.1), and the process was repeated with scVI tools. The obtained embeddings were then used for finding clusters with “FindNeighbors” and “FindClusters” functions and further visualized with UMAP dimensionality reduction with “RunUMAP” function using default parameters in Seurat. The optimal clustering threshold was chosen as 0.2 based on visual inspection of the clustering results in the UMAP reduced space. The markers used to define the clusters were found with Student’s t-test using the “FindMarkers” function in Seurat with logfc.threshold = 0.25 from expression data that was scaled with “ScaleData” function with scaling factor of 10000. Patients C141, C142, and C144 have moderate COVID-19. Patients reported12 to have severe (C143 and C145) or critical disease (C146, C148, C149, and C152) were considered to

## Data and code availability

Source code for TCRconv and all data used for training and testing the model are available at github.com/emmijokinen/tcrconv. Alternatively, epitope-specific data is available at vdjdb.cdr3.net, (VDJdb data) and https://doi.org/10.21417/ADPT2020COVID (ImmuneCODE data). Implementation for the scRNA+TCR*αβ*-seq data analysis is available at https://github.com/janihuuh/tcrconv_manu. Repertoire data is available at https://doi.org/10.21417/ADPT2020COVID (ImmuneCODE COVID-19 repertoires), and https://doi.org/10.21417/B7001Z (control repertoires from Emerson et al.). Data from Liao et al. is available at GEO GSE145926.

## Acknowledgements

We would like to acknowledge the computational resources provided by the Aalto Science-IT.

## Author Contributions

The project was originally proposed by HL, VG and RB. EJ, AD, MH, VG, and HL contributed to the model design. EJ, JH, MH, and HL designed the analysis. EJ implemented the model and performed the analysis. AD implemented the computation of TCR embeddings. JH implemented the analysis with scRNA-TCR*αβ*-seq data. MH, SM, and HL supervised the study. EJ, AD, MH, JH, and HL wrote the article. All authors contributed to the final version of the article.

