## Supplementary information for "Predicting recognition between T cell receptors and epitopes using contextualized motifs"

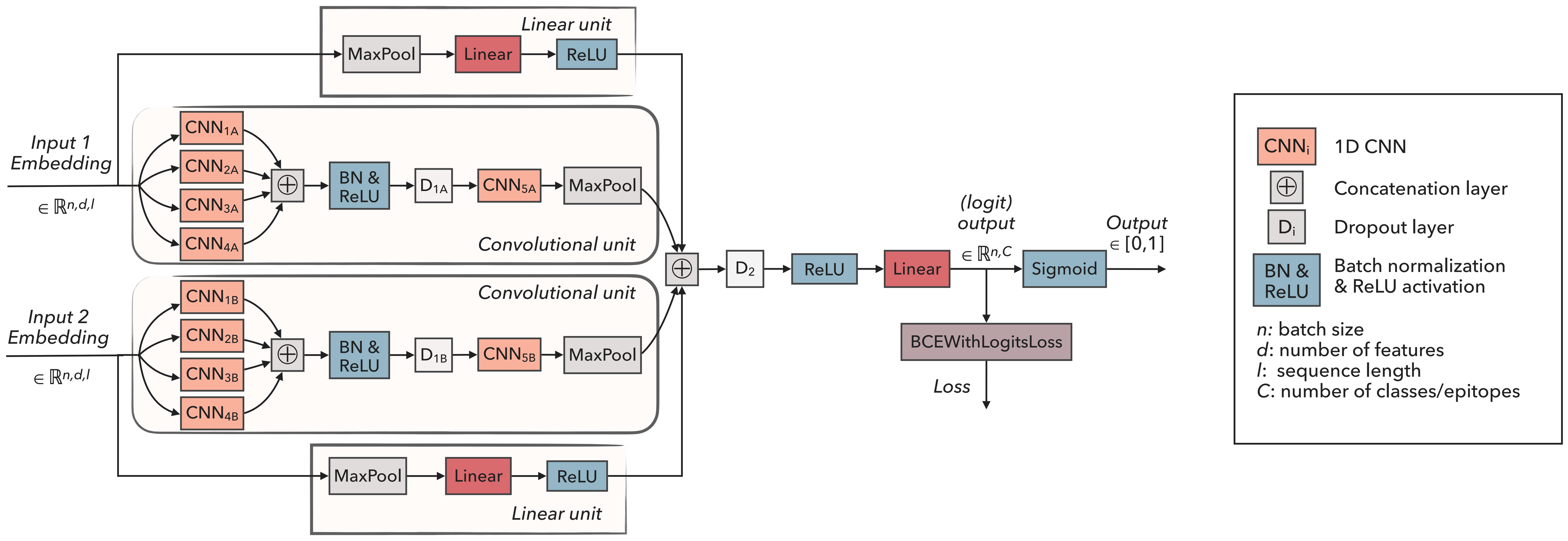


Supplementary Fig. 1. TCRconv multilabel predictor. TCRconv can utilize one or two inputs, i.e. embedding for TCRβ and/or TCRα. Each embedding goes through a convolutional and a linear unit in parallel. The outputs from these units (for each TCR chain) are concatenated and go through a final linear layer. During training, weighted binary crossentropy with logits loss (BCEWithLogitsLoss which includes a sigmoid function) is utilized, and the predictions are squashed between 0 and 1 by a sigmoid layer, separately for each epitope.

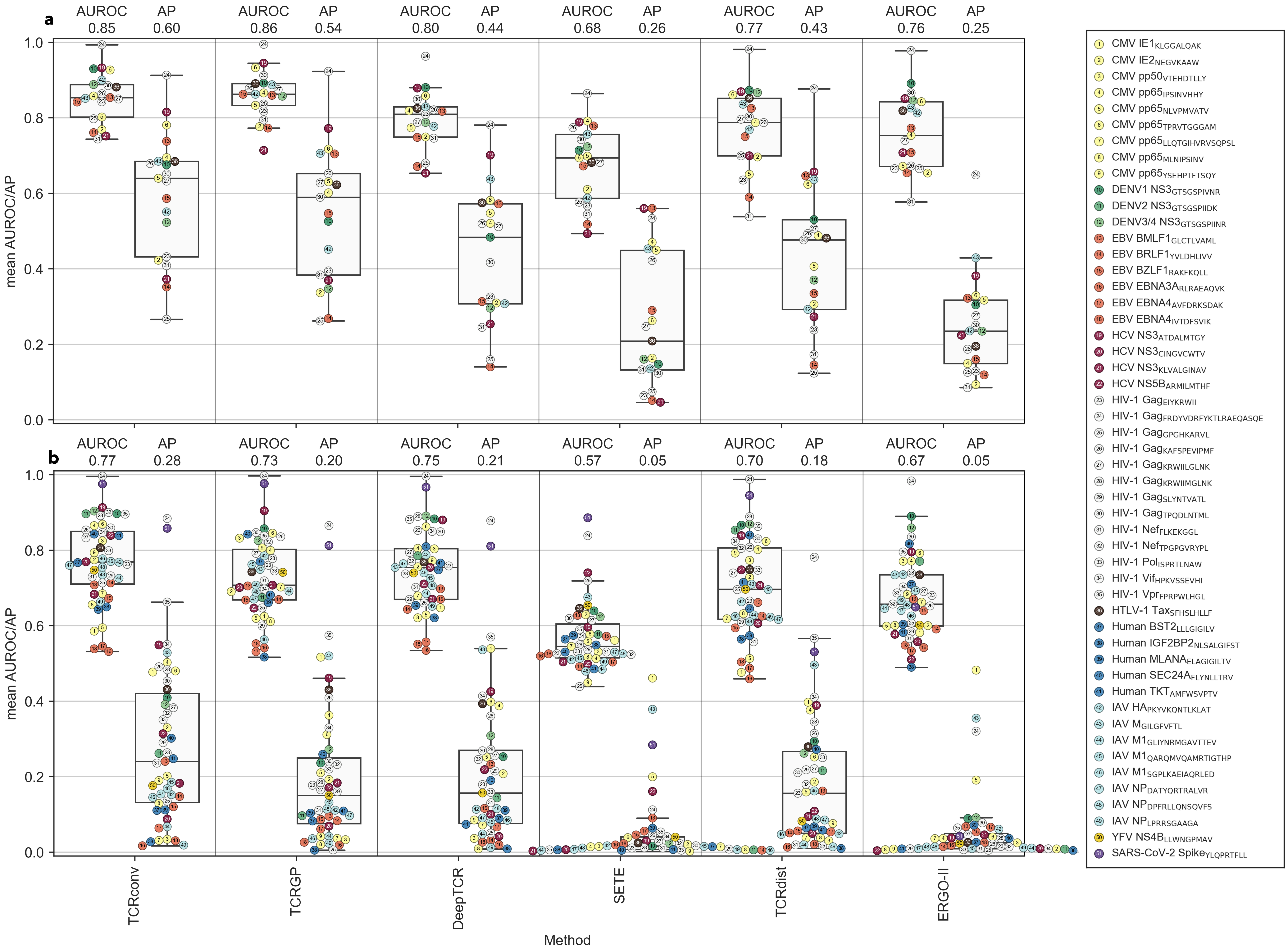


Supplementary Fig. 2. Method comparisons. Mean AUROC and AP scores on (a) VDJdbβ-small and (b) VDJdbβ-large dataset.

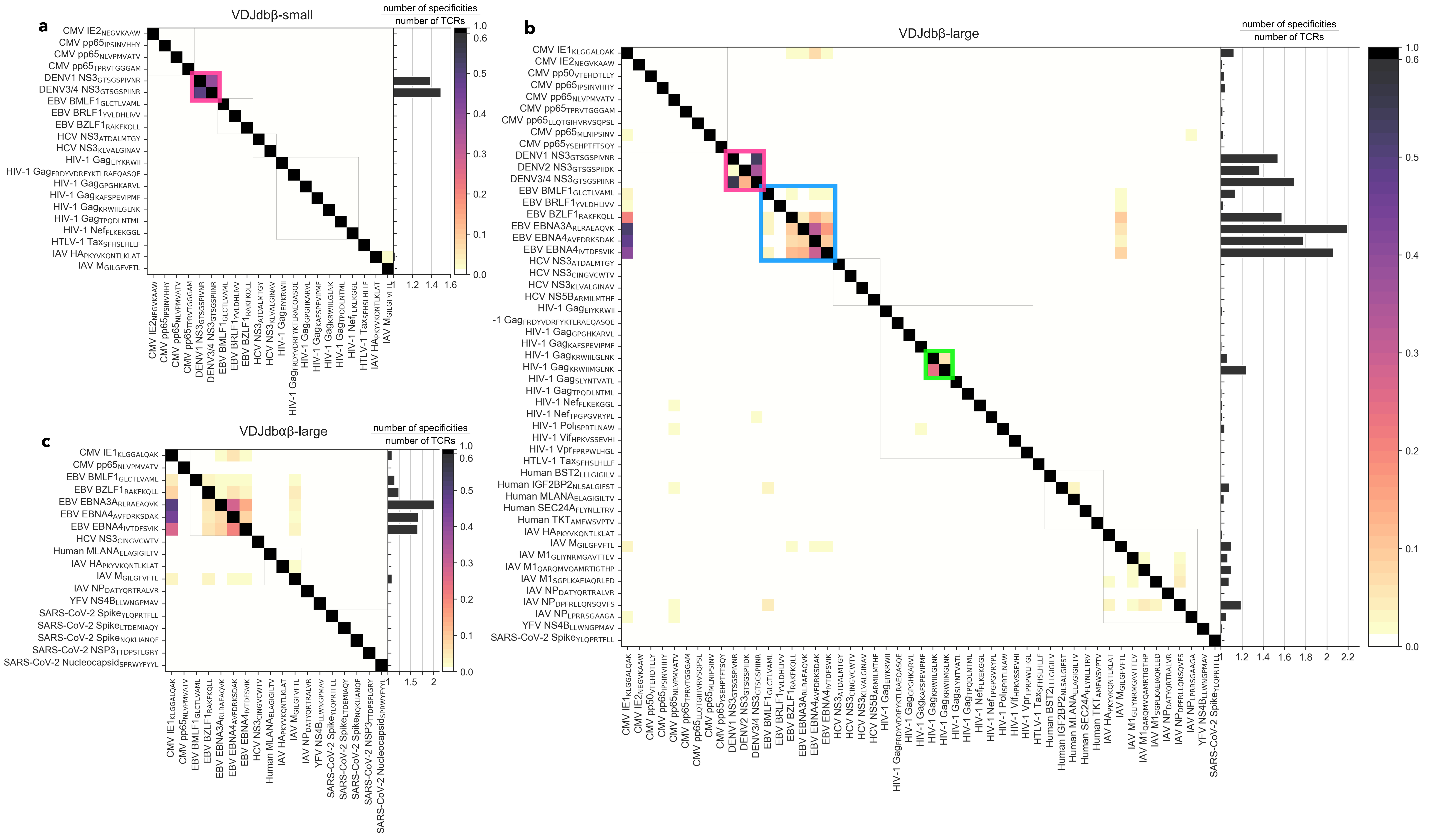


Supplementary Fig. 3. TCR cross-reactivity in datasets a) VDJdbβ-small, b) VDJdbβ-large, and c) VDJdbaβ-large. Each row of a heat map represents TCRs specific to the corresponding epitope and their fraction recognizing any of the epitopes present in the dataset. The bar plots on the right side of each heatmap show the average number of epitope specificities per TCR recognizing the epitope on the corresponding row. For example, TCRs specific to EBV epitope EBNA3A$_{RLRAEAQVK}$ recognize on average 2.2 different epitopes on (b) dataset VDJdbβ-large and 2.0 on (c) dataset VDJdbαβ-large. TCRs recognizing certain epitopes have notable cross-reactivity. To highlight them we have marked DENV epitopes with pink, EBV epitopes with blue, and two HIV-1 epitopes (HIV-1$_{KRWIILGLNK}$ and HIV-1$_{KRWIIMGLNK}$) with green.

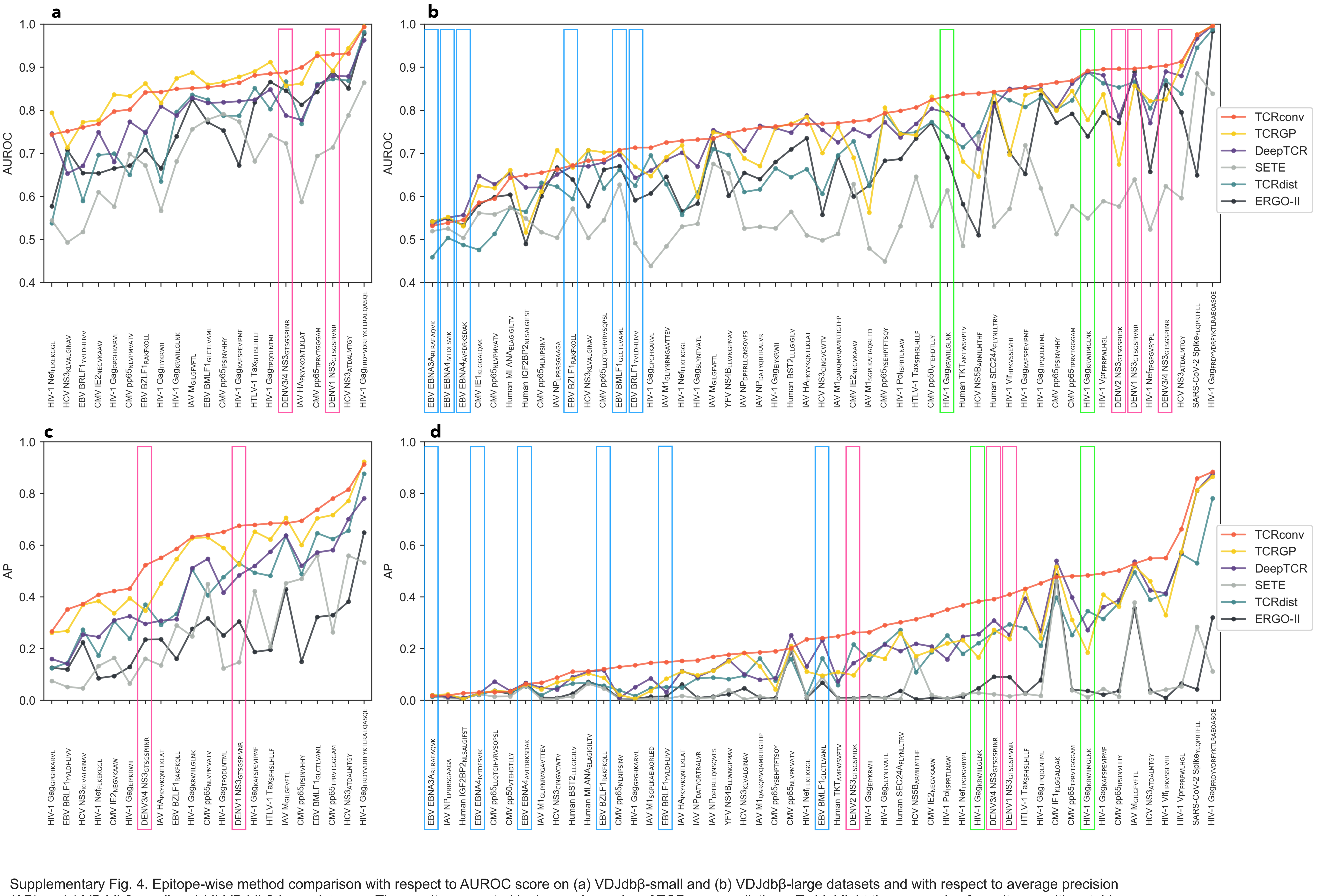


Supplementary Fig. 4. Epitope-wise method comparison with respect to AUROC score on (a) VDJdbβ-small and (b) VDJdbβ-large datasets and with respect to average precision (AP) on (c) VDJdbβ-small and (d) VDJdbβ-large datasets. The results are sorted by increasing order of TCRconv predictions. To highlight the accuracies for epitopes with notably cross-reactive TCRs, we have highlighted such epitopes similarly to Supplementary Fig. 3: DENV epitopes with pink, EBV epitopes with blue, and two HIV-1 epitopes (HIV-1$_{KRWIILGLNK}$ and HIV-1$_{KRWIIMGLNK}$) with green.

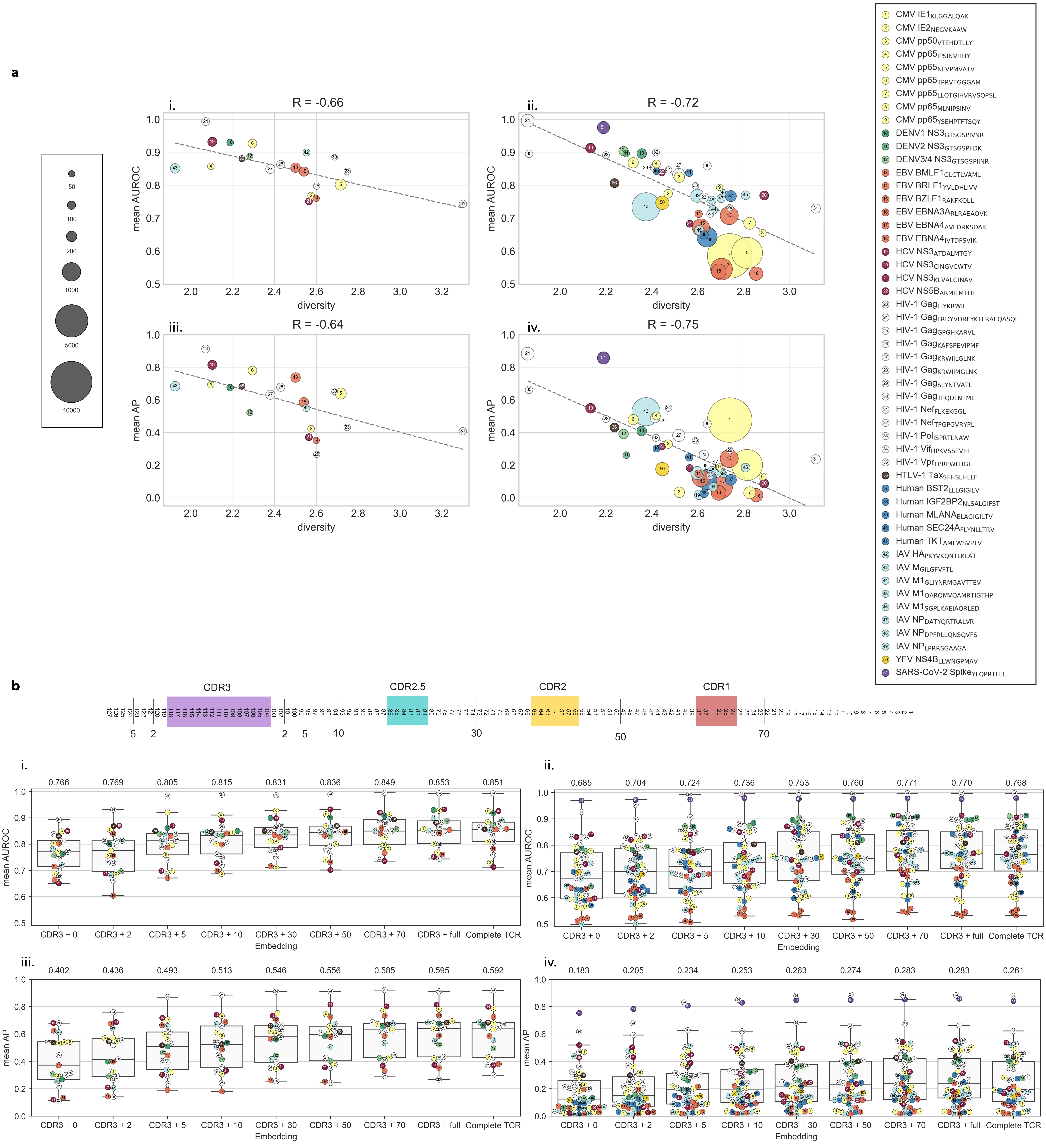


Supplementary Fig. 5. TCRconv evaluation. All AUROC and AP scores are obtained over stratified 10-fold cross-validation.
(a) Pearson's correlation between the diversity of epitope specific TCRs and the AUROC and AP scores. Panels (i) and (ii) show the mean AUROC scores for datasets VDJdbβ-small and VDJdbβ-large, respectively, and (iii) and (iv) mean AP scores for both datasets.
(b) Increasing embedding context size increases the predictive AUROC and AP scores. The schematics on the top show the approximate sections included in different context sizes. Complete TCR refers to using the complete TCR with the predictor, without extracting only the CDR3 part. Panels (i) and (ii) show the mean AUROC scores for datasets VDJdbβ-small and VDJdbβ-large, respectively, and (iii) and (iv) mean AP scores for both datasets.

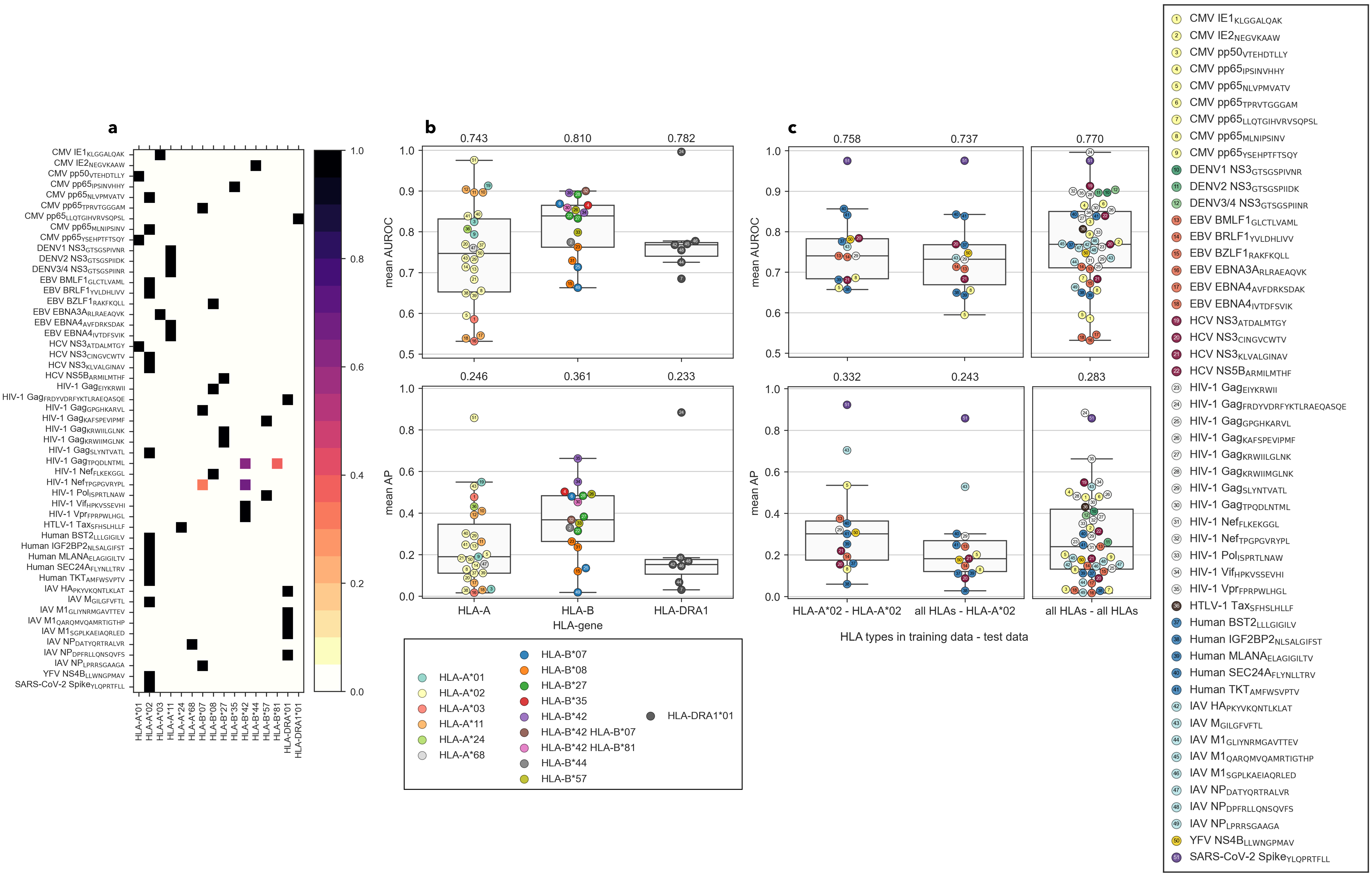


Supplementary Fig. 6. HLA-types of the MHCs restricting the epitopes do not alone explain variance in results. All AUROC and AP scores are obtained over stratified 10-fold cross-validation.

(a) HLA-types of the MHCs restricting the epitopes in dataset VDJdbβ-large.

(b) TCRconv predictions for VDJdbβ-large dataset have some variation in terms of AUROC and AP scores when the predictions are divided into three groups (HLA-A, HLA-B, and HLA-DRA1) based on the HLA-gene.

(c) AUROC and AP scores for HLA-A*02 restricted epitopes are similar whether the TCRconv is trained only on TCRs specific to HLA-A*02 restricted epitopes or to TCRs specific to all epitopes in VDJdbβ-large dataset. For reference TCRconv model trained and tested with TCRs specific to all epitopes, corresponding to results shown in Fig. 1a ii, is shown on right.

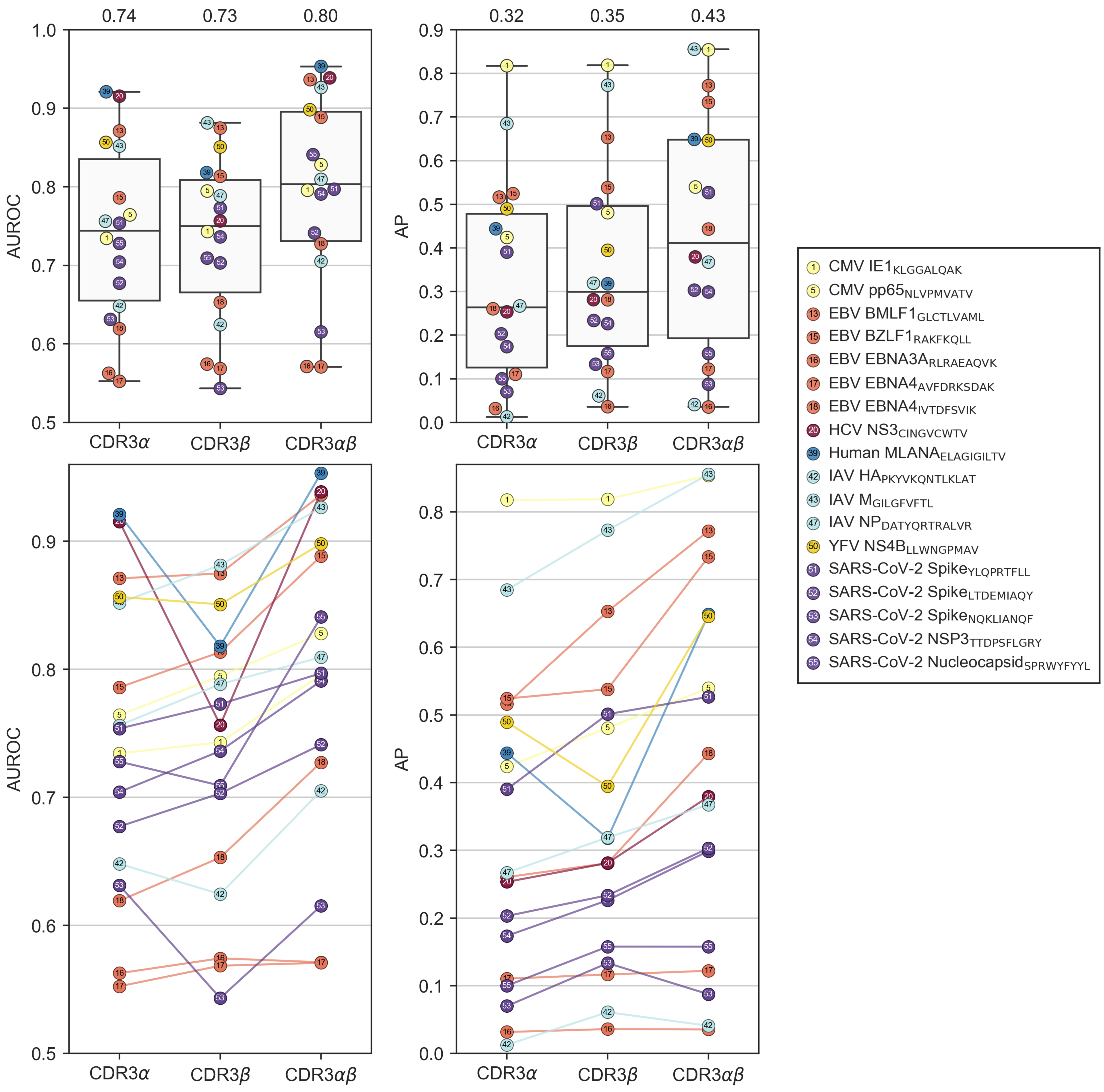


Supplementary Fig. 7. TCRconv performs best when using both α- and β-chains. Results are obtained on VDJdbaβ-large dataset in terms of average AUROC and AP scores over stratified 10-fold cross-validation. Each circle corresponds to TCRs specific to one epitope as described in the legend. Above boxplots show the distribution of the prediction accuracies when the TCRconv model is trained using embeddings for CDR3α, CDR3β, or both (always with the full context). Mean metrics are shown on top of each boxplot. Below the circles from the three models are connected by lines, illustrating how for most epitopes the best results are obtained when using both chains and using the β-chains is better than using α-chains, although there are exceptions.

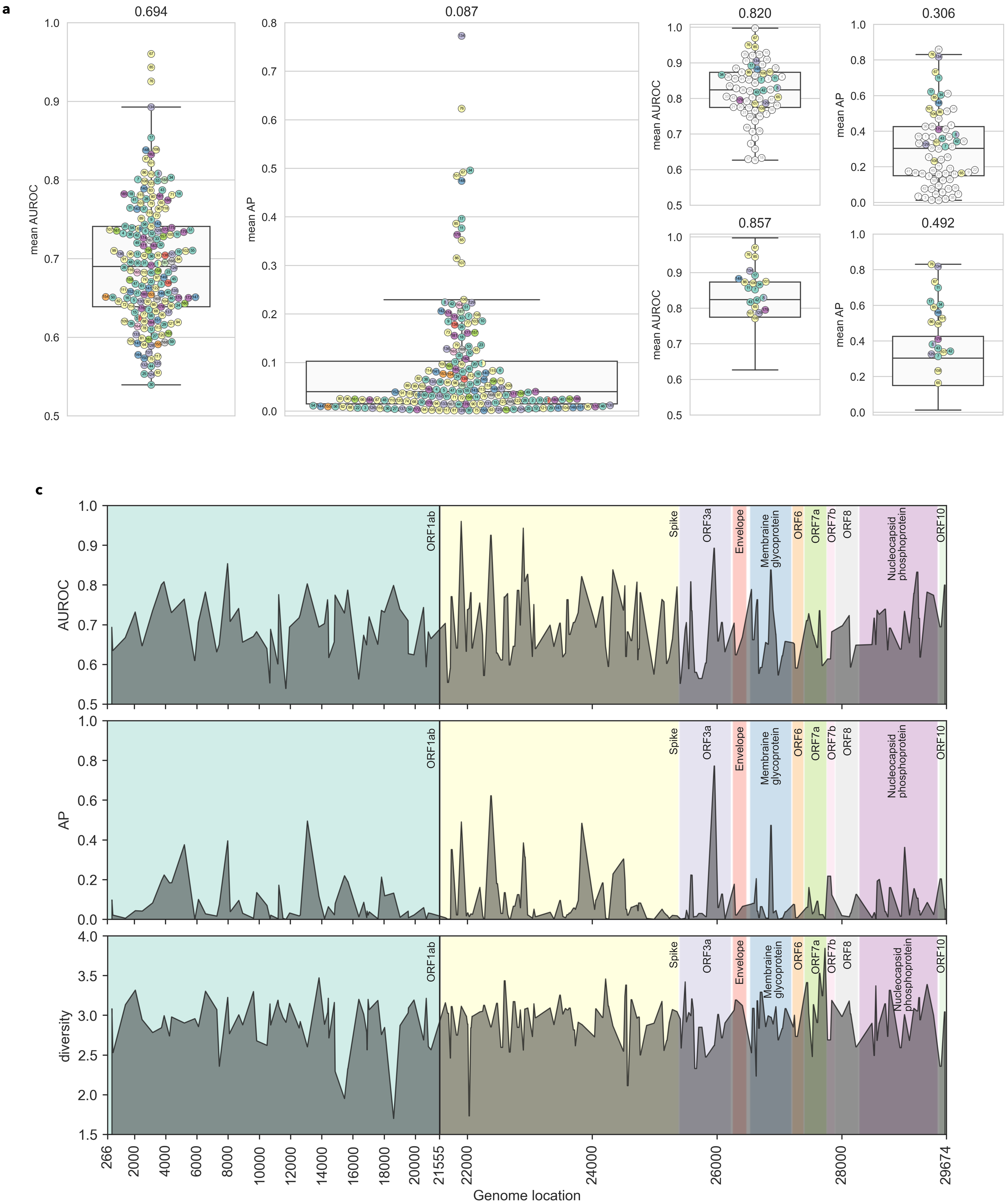


Supplementary Fig. 8. TCRconv prediction performance for SARS-CoV-2 epitopes.
(a) TCRconv performance in terms of AUROC and AP scores when trained with 139099 TCRs specific to 188 peptide groups from SARS-CoV-2. Mean scores are shown above both boxplots. Each circle represents the score for one peptide group, colored by the genomic region and numbered according to Supplementary Table 2.
(b) TCRconv performance when trained with TCRs specific to 20 best performing peptides groups from SARS-CoV-2 combined with VDJdbβ-large dataset; above results for all 70 peptide (groups) and below for only the 20 SARS-CoV-2 peptides. For SARS-CoV-2 peptides coloring and numbering are the same as in panel (a), other epitopes are white, and the numbering corresponds to Supplementary Table 1.
(c) AUROC and AP scores from the model from (a) by the peptides' genome location and the diversity of the TCRs specific to each peptide group by the peptides' genome location.

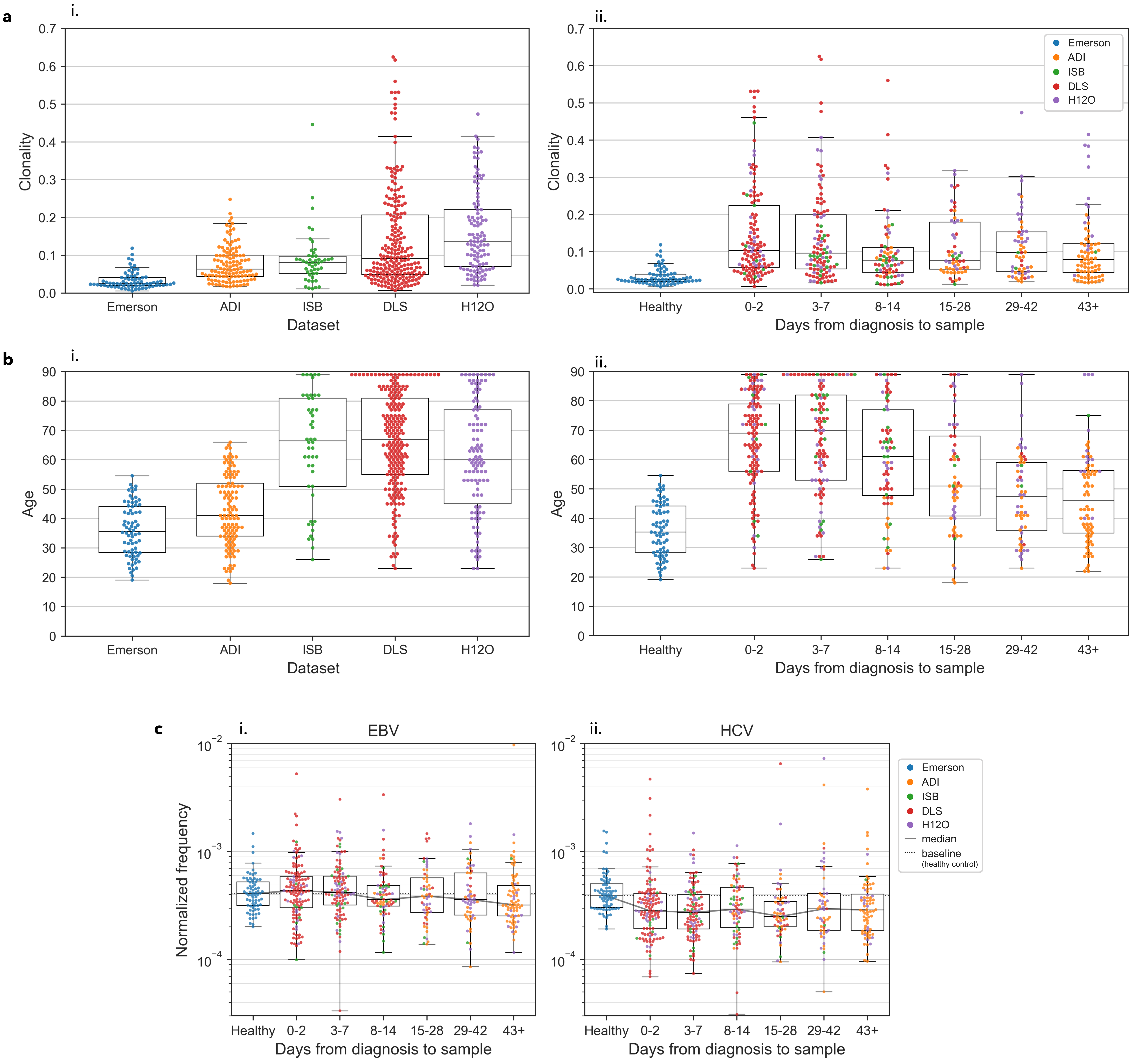


Supplementary Fig. 9. Analysis with COVID-19 patient repertoires.
(a) Shannon clonality (i) for each dataset and (ii) by Days from diagnosis to sample.
(b) Subject age (i) by dataset and (ii) by Days from diagnosis to sample.
(c) Normalized frequency grouped by number of days from diagnosis to sample for (i) six EBV specific epitopes and (ii) for four HCV specific epitopes.

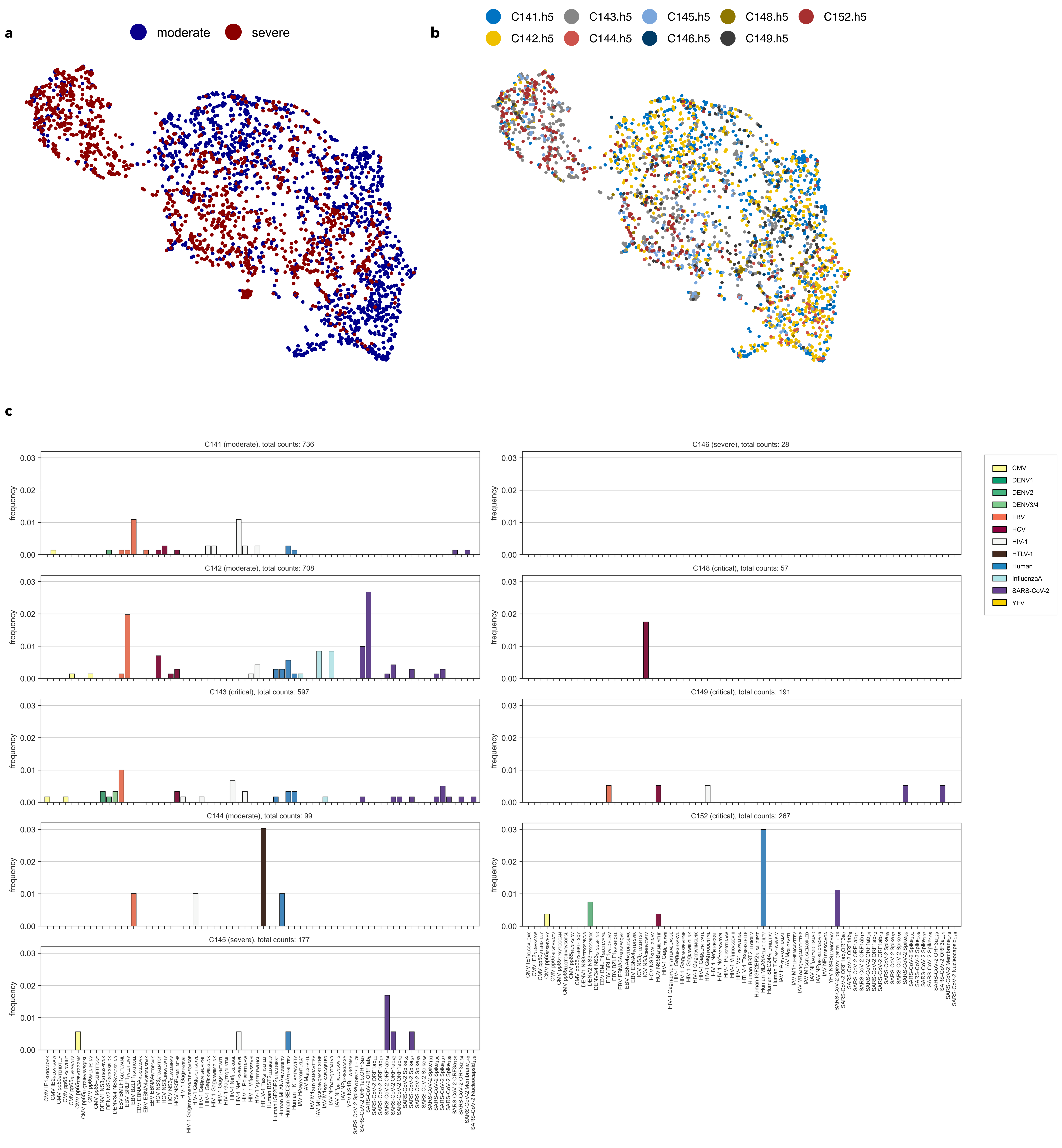


Supplementary Fig. 10. Characteristics of scRNA+TCR UMAP representation of CD8+ T-cells based on their phenotypes. Colored by (a) disease severity and by (b) patient. Patients C141, C142, and C144 have moderate COVID-19, while patients C143, C145, C146, C148, C149, and C152 have severe disease. (c) Frequencies of T-cells predicted to be specific to the tested epitopes separately for each patient. Only T-cells with both TCR- and RNA-seq available are shown.

Supplementary Table 1. Three datasets of epitope-specific TCR-data collected from VDJdb.The datasets contain epitope-specific TCRs for Cytomegalovirus (CMV), Dengue virus types 1, 2 and 3 (DENV1, DENV2, DENV3-4), Epstein-Barr virus (EBV), Hepatitis C virus (HCV), Human immunodeficiency virus type 1 (HIV-1), Influenza A virus (IAV), Severe acute respiratory syndrome coronavirus 2 (SARS-CoV-2), and Yellow Fever virus (YFV), as well as human stromal antigen 2 (BST2), insulin like growth factor 2 mRNA binding protein 2 (IGF2BP2), melanoma antigen (MLANA), and transketolase (TKT).

| # | Epitope Species | Epitope Gene | Epitope | MHC chain 1 | MHC chain 2 | VDJdbβ -large | VDJdbβ -small | VDJdbαβ -large |
|---|---|---|---|---|---|---|---|---|
| 1 | CMV | IE1 | KLGGALQAK | HLA-A*03 | B2M | 12693 | | 13664 |
| 2 | CMV | IE2 | NEGVKAAW | HLA-B*44 | B2M | 118 | 62 | |
| 3 | CMV | pp50 | VTEHDTLLY | HLA-A*01 | B2M | 202 | | |
| 4 | CMV | pp65 | IPSINVHHY | HLA-B*35 | B2M | 92 | 58 | |
| 5 | CMV | pp65 | NLVPMVATV | HLA-A*02 | B2M | 4488 | 244 | 175 |
| 6 | CMV | pp65 | TPRVTGGGAM | HLA-B*07 | B2M | 197 | 122 | |
| 7 | CMV | pp65 | LLQTGIHVRVSQPSL | HLA-DRA1*01 | HLA-DRB1*15 | 304 | | |
| 8 | CMV | pp65 | MLNIPSINV | HLA-A*02 | B2M | 73 | | |
| 9 | CMV | pp65 | YSEHPTFTSQY | HLA-A*01 | B2M | 52 | | |
| 10 | DENV1 | NS3 | GTSGSPIVNR | HLA-A*11 | B2M | 165 | 59 | |
| 11 | DENV2 | NS3 | GTSGSPIIDK | HLA-A*11 | B2M | 60 | | |
| 12 | DENV3-4 | NS3 | GTSGSPIINR | HLA-A*11 | B2M | 158 | 46 | |
| 13 | EBV | BMLF1 | GLCTLVAML | HLA-A*02 | B2M | 969 | 159 | 279 |
| 14 | EBV | BRLF1 | YVLDHLIVV | HLA-A*02 | B2M | 79 | 51 | |
| 15 | EBV | BZLF1 | RAKFKQLL | HLA-B*08 | B2M | 842 | 151 | 1212 |
| 16 | EBV | EBNA3A | RLRAEAQVK | HLA-A*03 | B2M | 410 | | 422 |
| 17 | EBV | EBNA4 | AVFDRKSDAK | HLA-A*11 | B2M | 1642 | | 1723 |
| 18 | EBV | EBNA4 | IVTDFSVIK | HLA-A*11 | B2M | 550 | | 713 |
| 19 | HCV | NS3 | ATDALMTGY | HLA-A*01 | B2M | 169 | 139 | |
| 20 | HCV | NS3 | CINGVCWTV | HLA-A*02 | B2M | 131 | | 76 |
| 21 | HCV | NS3 | KLVALGINAV | HLA-A*02 | B2M | 65 | 65 | |
| 22 | HCV | NS5B | ARMILMTHF | HLA-B*27 | B2M | 66 | | |
| 23 | HIV-1 | Gag | EIYKRWII | HLA-B*08 | B2M | 148 | 60 | |
| 24 | HIV-1 | Gag | FRDYVDRFYKTLRAEQASQE | HLA-DRA*01 | HLA-DRB1*01,07,11,15, HLA-DRB5*01 | 367 | 95 | |
| 25 | HIV-1 | Gag | GPGHKARVL | HLA-B*07 | B2M | 66 | 53 | |
| 26 | HIV-1 | Gag | KAFSPEVIPMF | HLA-B*57 | B2M | 175 | 104 | |
| 27 | HIV-1 | Gag | KRWIILGLNK | HLA-B*27 | B2M | 320 | 141 | |
| 28 | HIV-1 | Gag | KRWIIMGLNK | HLA-B*27 | B2M | 66 | | |
| 29 | HIV-1 | Gag | SLYNTVATL | HLA-A*02 | B2M | 57 | | |
| 30 | HIV-1 | Gag | TPQDLNTML | HLA-B*42,81 | B2M | 101 | 40 | |
| 31 | HIV-1 | Nef | FLKEKGGL | HLA-B*08 | B2M | 144 | 78 | |
| 32 | HIV-1 | Nef | TPGPGVRYPL | HLA-B*07,42 | B2M | 67 | | |
| 33 | HIV-1 | Pol | ISPRTLNAW | HLA-B*57 | B2M | 54 | | |
| 34 | HIV-1 | Vif | HPKVSSEVHI | HLA-B*42 | B2M | 54 | | |
| 35 | HIV-1 | Vpr | FPRPWLHGL | HLA-B*42 | B2M | 83 | | |
| 36 | HTLV-1 | Tax | SFHSLHLLF | HLA-A*24 | B2M | 132 | 45 | |
| 37 | Human | BST2 | LLLGIGILV | HLA-A*02 | B2M | 233 | | |
| 38 | Human | IGF2BP2 | NLSALGIFST | HLA-A*03 | B2M | 111 | | |
| 39 | Human | MLANA | ELAGIGILTV | HLA-A*02 | B2M | 1305 | | 388 |
| 40 | Human | SEC24A | FLYNLLTRV | HLA-A*02 | B2M | 61 | | |
| 41 | Human | TKT | AMFWSVPTV | HLA-A*02 | B2M | 82 | | |
| 42 | IAV | HA | PKYVKQNTLKLAT | HLA-DRA*01 | HLA-DRB1*01,04 | 388 | 69 | 59 |
| 43 | IAV | M1 | GILGFVFTL | HLA-A*02 | B2M | 3430 | 160 | 1815 |
| 44 | IAV | M1 | GLIYNRMGAVTTEV | HLA-DRA*01 | HLA-DRB1*01 | 121 | | |
| 45 | IAV | M1 | QARQMVQAMRTIGTHP | HLA-DRA*01 | HLA-DRB1*01 | 124 | | |
| 46 | IAV | M1 | SGPLKAEIAQRLED | HLA-DRA*01 | HLA-DRB1*01 | 64 | | |
| 47 | IAV | NP | DATYQRTRALVR | HLA-A*68 | B2M | 102 | | 92 |
| 48 | IAV | NP | DPFRLLQNSQVFS | HLA-DRA*01 | HLA-DRB1*01 | 104 | | |
| 49 | IAV | NP | LPRRSGAAGA | HLA-B*07 | B2M | 159 | | |
| 50 | YFV | NS4B | LLWNGPMAV | HLA-A*02 | B2M | 409 | | 239 |
| 51 | SARS-CoV-2 | Spike | YLQPRTFLL | HLA-A*02 | B2M | 315 | | 261 |
| 52 | SARS-CoV-2 | Spike | LTDEMIAQY | HLA-A*01 | B2M | | | 122 |
| 53 | SARS-CoV-2 | Spike | NQKLIANQF | HLA-B*15 | B2M | | | 71 |
| 54 | SARS-CoV-2 | NSP3 | TTDPSFLGRY | HLA-A*01 | B2M | | | 243 |
| 55 | SARS-CoV-2 | Nucleocapsid | SPRWYFYYL | HLA-B*07 | B2M | | | 75 |
| | | | | | TOTAL epitope-TCR pairs: | 32367 | 2001 | 21629 |
| | | | | | TOTAL unique TCRs | 30503 | 1977 | 20200 |

Supplementary Table 2. Method comparison. Mean AUROC and AP scores for TCRconv, TCRGP, TCRdist, SETE, DeepTCR and ERGO-II from stratified 10-fold cross-validation. Mean AUROC and AP scores are macro averages over all epitopes. Standard deviation is given over all folds and over all epitopes (Epit.), showing that with all methods variation between folds is smaller than variation between different epitopes. For TCRGP, DeepTCR and TCRdist the results were computed with models using only CDR3βs or additionally other components of TCRβs. With these methods accuracies were higher when additional components were used. All result figures present the more accurate version of each method.

| | | **VDJdbβ-small** | | | | | | **VDJdbβ-large** | | | | | |
|---|---|---|---|---|---|---|---|---|---|---|---|---|---|
| Method | TCRβ parts | Mean AUROC | Standard deviation Folds | Standard deviation Epit. | Mean AP | Standard deviation Folds | Standard deviation Epit. | Mean AUROC | Standard deviation Folds | Standard deviation Epit. | Mean AP | Standard deviation Folds | Standard deviation Epit. |
| TCRconv | CDR3 + full context | 0.853 | 0.028 | 0.064 | **0.595** | 0.054 | 0.164 | **0.770** | 0.010 | 0.108 | **0.283** | 0.016 | 0.205 |
| TCRGP | CDR3 | 0.801 | 0.029 | 0.074 | 0.451 | 0.045 | 0.195 | 0.675 | 0.010 | 0.117 | 0.168 | 0.011 | 0.177 |
| | all CDRs | **0.860** | 0.026 | 0.062 | 0.544 | 0.054 | 0.175 | 0.728 | 0.014 | 0.106 | 0.202 | 0.014 | 0.192 |
| DeepTCR | CDR3 | 0.752 | 0.019 | 0.088 | 0.356 | 0.024 | 0.188 | 0.705 | 0.007 | 0.108 | 0.173 | 0.009 | 0.173 |
| | CDR3+V | 0.797 | 0.022 | 0.072 | 0.438 | 0.031 | 0.174 | 0.747 | 0.005 | 0.101 | 0.213 | 0.012 | 0.194 |
| SETE | CDR3 | 0.679 | 0.022 | 0.099 | 0.265 | 0.027 | 0.175 | 0.569 | 0.012 | 0.086 | 0.049 | 0.003 | 0.091 |
| TCRdist | CDR3 | 0.702 | 0.027 | 0.082 | 0.334 | 0.028 | 0.150 | 0.641 | 0.008 | 0.116 | 0.131 | 0.008 | 0.139 |
| | all CDRs | 0.770 | 0.029 | 0.105 | 0.432 | 0.034 | 0.190 | 0.704 | 0.008 | 0.125 | 0.183 | 0.009 | 0.169 |
| ERGO-II | CDR3+V | 0.761 | 0.022 | 0.099 | 0.248 | 0.026 | 0.130 | 0.673 | 0.028 | 0.102 | 0.053 | 0.005 | 0.091 |

$$\text{standard deviation (epit.)} = \sqrt{\frac{\sum_{\mathrm{e}=1}^{N_\mathrm{e}} (S_\mathrm{e} - S)^2}{N_\mathrm{e} - 1}}$$

$$\text{standard deviation (folds)} = \sqrt{\frac{\sum_{\mathrm{f}=1}^{N_\mathrm{f}} (S_\mathrm{f} - S)^2}{N_\mathrm{f} - 1}}$$

$$S_\mathrm{e} = \sum_{\mathrm{f}=1}^{N_\mathrm{f}} \frac{S_{\mathrm{e,f}}}{N_\mathrm{f}}, \quad S_\mathrm{f} = \sum_{\mathrm{e}=1}^{N_\mathrm{e}} \frac{S_{\mathrm{e,f}}}{N_\mathrm{e}}, \quad S = \sum_{\mathrm{e}=1}^{N_\mathrm{e}} \sum_{\mathrm{f}=1}^{N_\mathrm{f}} \frac{S_{\mathrm{e,f}}}{N_\mathrm{e} N_\mathrm{f}}$$

$N_\mathrm{e}$ is the number of epitopes (21 in VDJdbβ-small, 51 in VDJdbβ-large),
$N_\mathrm{f}$ is the number of folds (10),
$S_{\mathrm{e,f}}$ is the mean score (AUROC or AP) for epitope $\mathrm{e}$ in fold $\mathrm{f}$
$S_\mathrm{e}$ is the mean score for epitope $\mathrm{e}$ over all folds,
$S_\mathrm{f}$ is the mean score for fold $\mathrm{f}$ over all epitopes, and
$S$ is the mean score over all epitopes and folds.

Supplementary Table 3. Overview of ImmuneCODE MIRA data used for training TCRconv classifiers for SARS-CoV-2 specific epitopes. Spike refers to surface glycoprotein, membrane to membrane glycoprotein, and nucleocapsid to nucleocapsid phosphoprotein. The peptide groups are ordered by the start point of their genomic location (Loc). The coloring of the genomic regions matches Supplementary Fig. 8.

| # | ORF | Loc | Peptides | TCRs |
|---|---|---|---|---|
| 1 | ORF1ab, *spike* | 533 | AELEGIQY, *TLADAGFIK, LADAGFIKQY, ADAGFIKQY* | 108 |
| 2 | ORF1ab | 587 | VPHVGEIPVAY, GEIPVAYRKVLL | 709 |
| 3 | ORF1ab | 1391 | SEVGPEHSLAEY | 318 |
| 4 | ORF1ab | 2024 | TSDLATNNLVVMAY | 68 |
| 5 | ORF1ab | 2468 | APKEIIFL, KEIIFLEGETL | 2444 |
| 6 | ORF1ab | 3137 | KPLEFGATSAAL | 716 |
| **7** | ORF1ab | 3707 | LLSAGIFGA | 52 |
| **8** | ORF1ab,*ORF3a* | 3875 | AEIPKEEVKPF, *SASKIITLK, ASKIITLKK* | 170 |
| 9 | ORF1ab | 4211 | ALRKVPTDNYITTY, KVPTDNYITTY | 643 |
| 10 | ORF1ab | 4346 | SNEKQEILGTVSW, ILGTVSWNL | 632 |
| **11** | ORF1ab | 5171 | HTTDPSFLGRY | 10134 |
| 12 | ORF1ab | 5834 | SEYKGPITDVFY, ITDVFYKENSY | 376 |
| 13 | ORF1ab | 6074 | FADDLNQLTGY | 89 |
| 14 | ORF1ab | 6521 | ITEEVGHTDLMAAY | 226 |
| 15 | ORF1ab | 7253 | AYILFTRFFYV | 243 |
| 16 | ORF1ab | 7415 | YIFFASFYY | 457 |
| **17** | ORF1ab | 7952 | QLMCQPILL, QLMCQPILLL | 1062 |
| 18 | ORF1ab | 8060 | SSTFNVPMEKLK | 82 |
| 19 | ORF1ab | 8111 | AEAELAKNVSL, AELAKNVSLDNVL | 1866 |
| 20 | ORF1ab | 8660 | HTDFSSEIIGY | 56 |
| 21 | ORF1ab | 8915 | FLPRVFSAV | 1520 |
| 22 | ORF1ab | 9602 | TFYLTNDVSFL | 53 |
| 23 | ORF1ab | 9812 | FLLNKEMYL | 85 |
| 24 | ORF1ab | 10472 | FLNGSCGSV | 4506 |
| 25 | ORF1ab | 10664 | VLAWLYAAV | 263 |
| 26 | ORF1ab | 10709 | FLNRFTTTL | 235 |
| 27 | ORF1ab | 11168 | FLCLFLLPSL, FLLPSLATV | 339 |
| 28 | ORF1ab | 11228 | MPASWVMRI | 815 |
| 29 | ORF1ab | 11492 | NYSGVVTTVMF | 72 |
| 30 | ORF1ab | 11684 | TLGVYDYLV, GVYDYLVST | 104 |
| 31 | ORF1ab | 11921 | KLWAQCVQL | 741 |
| 32 | ORF1ab | 12530 | FTYASALWEI | 69 |
| 33 | ORF1ab | 12599 | SEISMDNSPNL | 208 |
| **34** | ORF1ab | 13058 | TVLSFCAFA, VLSFCAFAV | 1095 |
| 35 | ORF1ab | 13805 | YTMADLVYA, TMADLVYAL | 373 |
| 36 | ORF1ab | 14159 | LLMPILTLT | 122 |
| 37 | ORF1ab | 14360 | ILHCANFNV | 353 |
| 38 | ORF1ab | 14402 | FPPTSFGPL | 686 |
| 39 | ORF1ab | 14441 | FVDGVPFVV | 5402 |
| 40 | ORF1ab | 14789 | ISDYDYYRY | 71 |
| 41 | ORF1ab | 14840 | RQLLFVVEV | 1393 |
| **42** | ORF1ab | 15437 | MVMCGGSLYV, VMCGGSLYV | 879 |
| **43** | ORF1ab | 15641 | NRDVDTDFVNEFY, DTDFVNEFYAY | 326 |
| 44 | ORF1ab | 16358 | LVLSVNPYV | 110 |
| 45 | ORF1ab | 16673 | KLSYGIATV | 4607 |
| 46 | ORF1ab | 16862 | VVYRGTTTY | 222 |
| 47 | ORF1ab | 16889 | KLNVGDYFV | 372 |
| 48 | ORF1ab | 16952 | TLVPQEHYV | 274 |
| 49 | ORF1ab | 17024 | SSNVANYQK | 108 |
| 50 | ORF1ab | 17579 | IVDTVSALV | 124 |
| 51 | ORF1ab | 17810 | ILGLPTQTV | 451 |
| 52 | ORF1ab | 18011 | IPRRNVATL | 72 |
| 53 | ORF1ab | 18590 | VLWAHGFEL | 1392 |
| 54 | ORF1ab | 18998 | ALLADKFPV | 50 |
| 55 | ORF1ab | 19520 | YLDAYNMMI | 500 |
| 56 | ORF1ab | 19538 | MMISAGFSL | 51 |
| 57 | ORF1ab, [illegible] | 19895 | APAHISTI, *LIVNSVLLFL, SVLLFLAFV, LLFLAFVVFL* | 1555 |
| 58 | ORF1ab | 20510 | LLLDDFVEI, LLDDFVEII | 2031 |
| 59 | ORF1ab | 20681 | GVAMPNLYK | 51 |
| 60 | ORF1ab | 20816 | YLNTLTLAV | 828 |
| 61 | ORF1ab | 20987 | TLIGDCATV | 856 |
| 62 | spike | 21632 | LPPAYTNSF | 145 |
| 63 | spike | 21668 | VYYPDKVFR, YPDKVFRSS, YPDKVFRSSV, KVFRSSVLH | 121 |
| 64 | spike | 21710 | STQDLFLPFF, TQDLFLPFF | 155 |
| **65** | spike | 21725 | FLPFFSNVT, LPFFSNVTW, LPFFSNVTWF, PFFSNVTWF, FFSNVTWFH, SNVTWFHAI | 455 |
| 66 | spike | 21809 | VLPFNDGVY, VLPFNDGVYF, LPFNDGVYF, LPFNDGVYFA, DGVYFASTEK, GVYFASTEK | 1467 |
| **67** | spike | 21887 | TLDSKTQSL | 171 |
| 68 | spike | 21965 | FCNDPFLGVY, CNDPFLGVY, CNDPFLGVYY | 224 |
| 69 | spike, *membrane* | 21986 | GVYYHKNNK, YYHKNNKSW, *VPLHGTIL* | 62 |
| 70 | spike | 22010 | KSWMESEFRV, SWMESEFRVY, WMESEFRVY | 179 |
| 71 | spike | 22037 | VYSSANNCTF, SSANNCTFEY | 240 |
| 72 | spike | 22058 | CTFEYVSQPF, FEYVSQPFL, EYVSQPFLM | 427 |
| 73 | spike | 22184 | TPINLVRDL | 268 |
| 74 | spike | 22229 | LEPLVDLPI | 536 |
| 75 | spike | 22244 | DLPIGINITR, LPIGINITRF, INITRFQTL | 99 |
| **76** | spike | 22355 | YYVGYLQPRTF, YLQPRTFL, YLQPRTFLL | 1217 |
| 77 | spike | 22451 | SETKCTLKSF, CTLKSFTVEK, TLKSFTVEK | 150 |
| 78 | spike | 22520 | VQPTESIVRF, QPTESIVRF, TESIVRFPNI, VRFPNITNL, RFPNITNLCPF, FPNITNLCPF | 1460 |
| 79 | spike | 22574 | FGEVFNATRF, GEVFNATRF, FNATRFASVY, NATRFASVY | 836 |
| 80 | spike | 22631 | RISNCVADY | 67 |
| 81 | spike | 22655 | YSVLYNSASF, SVLYNSASF, LYNSASFSTF, NSASFSTFK | 62 |
| 82 | spike | 22718 | CFTNVYADSF, FTNVYADSF, FTNVYADSFV, KLNDLCFTNV, LNDLCFTNVY, NVYADSFVIR, VYADSFVIR | 484 |
| 83 | spike | 22784 | APGQTGKIA, GQTGKIADY, KIADYNYKL, QTGKIADYNY, RQIAPGQTGK | 118 |
| 84 | spike | 22832 | KLPDDFTGCV | 1381 |
| **85** | spike | 22880 | NLDSKVGGNY | 54 |
| **86** | spike | 22904 | NYLYRLFRK, NYNYLYRLF | 536 |
| 87 | spike | 22946 | FERDISTEI, FERDISTEIY, KPFERDISTEI | 223 |
| 88 | spike | 23027 | YFPLQSYGF | 434 |
| 89 | spike | 23051 | FQPTNGVGY | 62 |
| 90 | spike | 23072 | GYQPYRVVVL, PYRVVVLSF, QPYRVVVL, QPYRVVVLSF | 263 |
| 91 | spike | 23309 | EILDITPCSF | 278 |
| 92 | spike | 23372 | TSNQVAVLY | 87 |
| 93 | spike | 23465 | STGSNVFQTR, TGSNVFQTR, VYSTGSNVF | 348 |
| 94 | spike | 23513 | GAEHVNNSY, IGAEHVNNSY | 83 |
| 95 | spike | 23552 | IGAGICASY, IPIGAGICASY | 136 |
| 96 | spike | 23579 | QTNSPRRAR, SPRRARSVA, SYQTQTNSPR, TQTNSPRRAR | 115 |
| 97 | spike | 23615 | ASQSIIAYTM, RSVASQSII, SIIAYTMSL, SQSIIAYTM, VASQSIIAY | 721 |
| 98 | spike | 23678 | AIPTNFTISV, AYSNNSIAIPTNF, IPTNFTISV, NSIAIPTNF | 771 |
| 99 | spike | 23714 | FTISVTTEIL | 204 |
| 100 | spike | 23759 | KTSVDCTMYI | 174 |
| **101** | spike | 23816 | LLLQYGSFC, LLQYGSFCT | 93 |
| 102 | spike | 24023 | LLFNKVTLA | 101 |
| 103 | spike | 24140 | LLTDEMIAQY, LTDEMIAQY, LTDEMIAQYT, VLPPLLTDEMIAQY | 1080 |
| 104 | spike | 24200 | GTITSGWTF | 72 |
| 105 | spike | 24242 | IPFAMQMAY, LQIPFAMQM | 83 |
| **106** | spike | 24317 | NQKLIANQF | 82 |
| **107** | spike | 24485 | SVLNDILSR, VLNDILSRL | 55 |
| **108** | spike | 24509 | RLDKVEAEV | 50 |
| 109 | spike | 24521 | AEVQIDRLI, AEVQIDRLIT, VEAEVQIDRL, VQIDRLITGR | 246 |
| 110 | spike | 24548 | GRLQSLQTY, LITGRLQSL, RLQSLQTYV | 464 |
| 111 | spike | 24608 | AEIRASANL, AEIRASANLA, ASANLAATK | 211 |
| 112 | spike | 24701 | HLMSFPQSA, YHLMSFPQSA | 106 |
| 113 | spike | 24716 | FPQSAPHGV, FPQSAPHGVVF | 222 |
| 114 | spike | 24728 | APHGVVFL, APHGVVFLHV, GVVFLHVTY, VVFLHVTYV | 403 |
| 115 | spike | 24752 | HVTYVPAQEK, TYVPAQEKNF, VTYVPAQEK | 73 |
| 116 | spike | 24857 | GTHWFVTQR | 111 |
| 117 | spike | 24968 | TVYDPLQPELDSFK | 53 |
| 118 | spike | 25103 | KEIDRLNEV | 80 |
| 119 | spike | 25136 | NLNESLIDL | 203 |
| 120 | spike | 25178 | QYIKWPWYI, YEQYIKWPW, YEQYIKWPWY | 1751 |
| 121 | spike | 25220 | FIAGLIAIV | 323 |
| 122 | spike | 25268 | CMTSCCSCLK, MTSCCSCLK | 159 |
| 123 | spike | 25343 | SEPVLKGVKL | 149 |
| 124 | ORF3a | 25393 | DLFMRIFTI, MDLFMRIFTI | 114 |
| 125 | ORF3a | 25408 | RIFTIGTVTLK | 140 |
| 126 | ORF3a | 25474 | FVRATATIPI | 53 |
| 127 | ORF3a | 25495 | IPIQASLPF | 124 |
| 128 | ORF3a | 25531 | IVGVALLAVF, VGVALLAVF | 54 |
| **129** | ORF3a | 25579 | ITLKKRWQL, LKKRWQLAL, TLKKRWQLA, TLKKRWQLAL | 143 |
| 130 | ORF3a | 25606 | ALSKGVHFV | 397 |
| 131 | ORF3a | 25627 | FVCNLLLLFV, LLFVTVYSHL, TVYSHLLLV | 1495 |
| 132 | ORF3a | 25690 | FLQSINFVR, FLQSINFVRI, FLYLYALVYF, GLEAPFLYLY, INFVRIIMR, LQSINFVRI, LQSINFVRII, QSINFVRII, SINFVRIIMR, VYFLQSINF, VYFLQSINFV, YFLQSINFVR, YLYALVYFL | 3298 |
| 133 | ORF3a | 25801 | LLYDANYFL, LLYDANYFLC, LYDANYFLCW, NPLLYDANY, PLLYDANYFL, YDANYFLCW | 1039 |
| 134 | ORF3a | 25930 | SEHDYQIGGYTEKW, YQIGGYTEK, YQIGGYTEKW | 3447 |
| 135 | ORF3a | 25996 | SYFTSDYYQL, VLHSYFTSDY, YFTSDYYQLY | 1174 |
| 136 | ORF3a | 26050 | GVEHVTFFIY, HVTFFIYNK, STDTGVEHVTFFIY, VEHVTFFIY | 304 |
| 137 | ORF3a | 26113 | EEHVQIHTI | 178 |
| 138 | envelope | 26260 | SEETGTLIV | 63 |
| 139 | envelope | 26392 | SLVKPSFYV | 128 |
| 140 | membrane | 26538 | GTITVEELK | 62 |
| 141 | membrane | 26553 | EELKKLLEQW, KLLEQWNLV, QWNLVIGFLF | 187 |
| 142 | membrane | 26613 | WICLLQFAY | 595 |
| 143 | membrane | 26625 | FAYANRNRF, LQFAYANRNR, YANRNRFLY | 416 |
| 144 | membrane | 26652 | FLYIIKLIFL, FLYIIKLVFL, LYIIKLIFL, LYIIKLIFLW, RFLYIIKLIF, YIIKLIFLW, YIIKLIFLWL | 950 |
| 145 | membrane | 26679 | FLWLLWPVT, FLWLLWPVTL, LWLLWPVTL, LWPVTLACF, TLACFVLAAV, WLLWPVTLA, [illegible] | 5441 |
| 146 | membrane | 26763 | AIAMACLVGL, IAMACLVGLM | 112 |
| 147 | membrane | 26802 | FIASFRLFA, SYFIASFRLF, YFIASFRLF, YFIASFRLFA | 372 |
| 148 | membrane | 26844 | MWSFNPETNI, SFNPETNIL, SMWSFNPET | 412 |
| 149 | membrane | 26928 | SELVIGAVI, SELVIGAVIL | 1564 |
| 150 | membrane | 26964 | HLRIAGHHL, RIAGHHLGR | 101 |
| 151 | membrane | 27027 | ATSRTLSYY, ATSRTLSYYK, TSRTLSYYK, TVATSRTLSY | 334 |
| 152 | membrane | 27069 | ASQRVAGDSGFAAY, DSGFAAYSR, VAGDSGFAAY | 57 |
| 153 | ORF6 | 27208 | HLVDFQVTI | 174 |
| 154 | ORF6 | 27226 | VTIAEILLI | 130 |
| 155 | ORF6 | 27241 | FKVSIWNLDY, ILLIIMRTFK, IMRTFKVSI, KVSIWNLDY, KVSIWNLDYI, LIIMRTFKV, RTFKVSIWNL, SIWNLDYII, TFKVSIWNL, VSIWNLDYII | 727 |
| 156 | ORF7a | 27418 | CELYHYQECV, ITLATCELY, LITLATCELY, TLATCELYHY | 609 |
| 157 | ORF7a | 27454 | QECVRGTTVL | 203 |
| 158 | ORF7a | 27511 | YEGNSPFHPL | 101 |
| 159 | ORF7a | 27523 | HPLADNKFAL, SPFHPLADNKFAL | 252 |
| 160 | ORF7a | 27580 | CPDGVKHVY, DGVKHVYQL, FAFACPDGVKHVY | 187 |
| 161 | ORF7a | 27625 | RARSVSPKL, SVSPKLFIR | 136 |
| 162 | ORF7a | 27670 | ELYSPIFLI, LYSPIFLIV, QELYSPIFL, VQELYSPIF, VQELYSPIFL | 2332 |
| 163 | ORF7a | 27715 | VFITLCFTL, VFITLCFTLK | 138 |
| 164 | ORF7b | 27756 | AFLLFLVLI, FLAFLLFLV, FYLCFLAFL, FYLCFLAFLL, IDFYLCFLAF, IELSLIDFYL, LIDFYLCFL, LLFLVLIML, MIELSLIDFY, SLIDFYLCFL, YLCFLAFLL | 16778 |
| 165 | ORF7b | 27822 | IMLIIFWFSL, MLIIFWFSL | 1452 |
| 166 | ORF8 | 27984 | VDDPCPIHFY, VVDDPCPIHFY, YVVDDPCPI | 356 |
| 167 | ORF8 | 28104 | IQYIDIGNY | 241 |
| 168 | ORF8 | 28122 | GNYTVSCLPF, NYTVSCLPF, YTVSCLPFTI | 360 |
| 169 | ORF8 | 28206 | DFLEYHDVR, EDFLEYHDVR, LEYHDVRVV, LEYHDVRVVL, YEDFLEYHDVRVVL | 1749 |
| 170 | nucleocapsid | 28466 | FPRGQGVPI, KFPRGQGVPI | 62 |
| 171 | nucleocapsid | 28496 | NSSPDDQIGY, NTNSSPDDQIGYY, SSPDDQIGY, SSPDDQIGYY | 166 |
| 172 | nucleocapsid | 28529 | YYRRATRRIR | 84 |
| 173 | nucleocapsid | 28550 | RIRGGDGKM, RIRGGDGKMK | 107 |
| 174 | nucleocapsid | 28583 | LSPRWYFYY, SPRWYFYYL | 3084 |
| 175 | nucleocapsid | 28673 | ALNTPKDHI, ATEGALNTPK | 509 |
| 176 | nucleocapsid | 28718 | NPANNAAIV, NPANNAAIVL, RNPANNAAIV | 357 |
| 177 | nucleocapsid | 28745 | GTTLPKGFY, LQLPQGTTL, QLPQGTTLPK, TTLPKGFYA, VLQLPQGTTL | 129 |
| 178 | nucleocapsid | 28916 | ALALLLLDR, GDAALALLLL, LALLLLDRL, LLLDRLNQL, LLLLDRLNQ, LLLLDRLNQL | 870 |
| 179 | nucleocapsid | 28988 | QQQGQTVTK, QQQQGQTVTK | 80 |
| 180 | nucleocapsid | 29069 | KAYNVTQAF | 1450 |
| 181 | nucleocapsid | 29138 | ELIRQGTDY, QELIRQGTDY, QELIRQGTDYKHW | 166 |
| 182 | nucleocapsid | 29186 | APSASAFFGM, AQFAPSASA, ASAFFGMSR, SASAFFGMSR | 494 |
| 183 | nucleocapsid | 29219 | GMSRIGMEV, SRIGMEVTPSGTW | 55 |
| 184 | nucleocapsid | 29234 | GMEVTPSGTWL, MEVTPSGTWL, TPSGTWLTY, VTPSGTWLTY | 2003 |
| 185 | nucleocapsid | 29348 | AYKTFPPTEPK, KTFPPTEPK | 1645 |
| 186 | nucleocapsid | 29456 | LPAADLDDF | 138 |
| 187 | ORF10 | 29558 | AFPFTIYSL, GYINVFAFPF, INVFAFPFTI, MGYINVFAF, NVFAFPFTI, NVFAFPFTIY, YINVFAFPF | 5160 |
| 188 | ORF10 | 29630 | AQVDVVNFNL, NYIAQVDVV | 119 |

Supplementary Table 4. Healthy control and ImmuneCODE repertoire data used in the analysis for T-cell dynamics during COVID-19 (Fig. 2a). The controls consist of the first 72 TCR repertoires from healthy (CMV-) subjects in cohort 1 in the study of Emerson et al. that had over 250 000 TCRs, number of templates reported, and where the subject is known to be at least 18 years old (which is the age of the youngest subject in the ImmuneCODE data used here). From ImmuneCODE 493 repertoires with over 250 000 TCRs and "Days from diagnosis to sample" reported were selected from four separate datasets.

| Cohort type | Cohort name | Institution | Study description | Mean age and s.d. (years) | Number of samples | Samples with ≥ 250000 TCRs and Days from |
|---|---|---|---|---|---|---|
| Healthy control | Emerson | Fred Hutchinson Cancer Research Center | Human peripheral blood samples were obtained from the institution's Research Cell Bank biorepository of healthy bone marrow donors. Donors underwent CMV serostatus testing at the time the samples were taken | 36.1 ± 8.9 | | 72 |
| COVID-19 | ImmuneRACE (ADI) | Adaptive Biotechnologies | Whole blood samples were collected from subjects from 24 geographic areas in the US with active infection, in convalescent phase, or exposed to SARS-CoV-2 | 42.6 ± 11.9 | 123 | 118 |
| COVID-19 | ISB | Institute for Systems Biology | Whole blood samples collected under the INCOVE project at Providence St. Joseph Health (Seattle, WA). Subjects were enrolled during the active phase and monitored through disease. | 66.1 ± 16.7 | 157 | 48 |
| COVID-19 | DLS | Discovery Life Sciences | Whole blood samples collected during routine care in accute and convalescdent phases procured through Discovery Life Sciences (Huntsville, AL). | 64.1 ± 18.5 | 431 | 216 |
| COVID-19 | H12O | Hospital Universitario 12 de Octubre | Whole blood samples were collected at the Hospital Universitario 12 de Octubre (Madrid, Spain) during the active or convalescent phase. | 60.5 ± 19.1 | 612 | 111 |
| | | | | | TOTAL: | 110 + 493 |

Supplementary Table 5. Significance of case-control and age effects on frequency of virus specific T-cells. Linear regression analysis was performed to assess if COVID patients have significantly higher frequency of virus specific T-cells than healthy control subjects, and if frequencies are positively correlated with subjects' age (see Methods). **(a)** The Benjamini-Hochberg adjusted p-values representing the significance of $b_{cc} > 0$. **(b)** The Benjamini-Hochberg adjusted p-values representing the significance of $b_{age} > 0$. One-tailed t-test was used for computing the p-values and the multiple testing adjustments are done for each virus (column) separately. Adjusted p-values smaller than 0.1 are bolded.

**a)**

| Time interval | SARS-CoV-2 | CMV | IAV | EBV | HCV |
|---|---|---|---|---|---|
| 0-2 | 0.3443 | 0.9181 | 1.0000 | 1.0000 | 1.0000 |
| 3-7 | **0.0087** | 0.8991 | 1.0000 | 0.7298 | 0.9982 |
| 8-14 | **0.0596** | 1.0000 | 1.0000 | 0.7324 | 1.0000 |
| 15-28 | **0.0104** | 0.8760 | 1.0000 | 0.7707 | 1.0000 |
| 29-42 | 0.2457 | 1.0000 | 1.0000 | 0.6010 | 1.0000 |
| 43 | 0.1790 | 0.9486 | 1.0000 | 0.7892 | 1.0000 |

**b)**

| Time interval | SARS-CoV-2 | CMV | IAV | EBV | HCV |
|---|---|---|---|---|---|
| 0-2 | **0.0033** | 0.0970 | 0.1009 | 0.4553 | 0.4698 |
| 3-7 | 0.2982 | **0.0261** | 0.1380 | **0.0632** | 0.4236 |
| 8-14 | 0.1846 | 0.1186 | 0.6596 | 0.6491 | 0.4310 |
| 15-28 | **0.0166** | **0.0029** | 0.1296 | **0.0460** | 0.2211 |
| 29-42 | **0.0000** | 0.1024 | 0.6254 | 0.3875 | 0.1598 |
| 43 | **0.0157** | 0.0354 | 0.6434 | 0.3731 | 0.4855 |
